# From shielding effect to hierarchical structures: a coarse-grained description of diversity

**DOI:** 10.64898/2026.08.03.742630

**Authors:** Ga Ching Lui, Sidhartha Goyal

## Abstract

The shared trade-off in microbial resource-utilization strategies has been proposed to resolve the paradox of the plankton, which states that the diversity observed in nature greatly exceeds the theoretically predicted upper bound that limits the number of coexisting species to the number of available resources. However, three important aspects remain unaddressed in this line of work. First, trade-offs have been quantified not only for phenotypes associated with alternative resource utilization, but also for other modes of microbial interaction. Second, in natural systems, not all taxa are subjected to the same trade-offs. Third, existing trade-off-based models do not explain the empirically observed clustering of taxa according to functional similarity. Here, we extend the trade-off-based framework to incorporate multiple types of resources. We assume that different subsets of taxa are constrained by distinct sets of trade-offs. Under this framework, our model predicts the emergence of clusters: while taxa with similar strategies can belong to the same cluster, each cluster is sustained by taxa with substantially different strategies. The resulting system supports high diversity with an effectively unlimited number of coexisting taxa, yet can still be described as a low-diversity community in which the number of coexisting functional clusters does not exceed the theoretical upper bound.

## I. INTRODUCTION

The paradox of plankton [1] points to the seeming contradiction between the rich diversity observed in nature Maharjan *et al*. [2] and the theoretical prediction given by the competitive exclusion principle [3][4], which gives at steady state an upper bound to the number of coexisting taxa that interact through changing the environment (e.g., consuming resources). In essence, it refers to the idea that if more taxa coexist compared to the number of abiotic factors present in the system, then the system is overdetermined [5], meaning that there are fewer variables than equations, which in general are not solvable. This has been a half-century-old problem since raised by Hutchinson [1], and different theoretical frameworks have been proposed over the years to resolve the paradox. A rather straightforward approach would be to introduce heterogeneity into the system; thus, the system no longer stays at some well-mixed steady states and is not bounded by the competitive exclusion principle. This line of approach includes spatial heterogeneity [6] and temporal fluctuations, either through external perturbations [7] or internal dynamics [8]. Furthermore, in well-mixed systems where there is no heterogeneity, the number of solvable variables in these models can be increased by modifying the bacterial growth rate such that interactions are dependent on the taxa abundances [9][10], thus bypassing the aforementioned upper bound.

Despite a long history of exploration, this problem remains an active research topic. Recent efforts consider various mechanisms in well-mixed systems that can potentially change the upper bound on the number of coexisting taxa. This includes increasing the dimensionality of the models by considering interactions, such as allelopathy [11] and cross-feeding [12][13][14][15][16], through secondary metabolites, which are by-products secreted during growth. In particular, cross-feeding has shown great success in accounting for experimental data under nutrient limitations [17][18]. The idea is to increase the upper bound by adding “hidden variables”, which are not measured nor monitored in real systems but can nonetheless influence the dynamics of the system. Another prominent idea is to eliminate the upper bound such that any arbitrary number of taxa can be accommodated through trade-offs in phenotypes. For competition over alternate resources, such as multiple sources of carbon, it has been shown in theoretical work [19][20][21][22] that the consortium of bacteria that shares the same trade-off can self-organize to flatten the fitness landscape, thereby sustaining a high diversity. In other words, the consortium maintains an internal environment that is shielded or decoupled from external changes [21]. We refer to this as the shielding effect.

Given these numerous proposed solutions to the paradox of plankton, why is it still of interest to us in today’s context? One of the relevant ideas that has not been fully explored is the connection of this problem with the coarse-grained description of diversity. Compared to the work that relies on hidden variables and models with high dimensionality such as in [16], coarse-graining points to a different line of thinking: diverse systems are governed by lower-dimensional dynamics that leads to clustering. In nature, hierarchical organizations of microbial communities have been observed in various systems, from soil and groundwater systems to the gut microbiome [23][24][25][26]. Theoretical work based on resource competition over alternate resources (e.g. different carbon sources) shows that taxa can be clustered based on their phenotypical similarities: the spectrum of the interaction matrix is largely dominated by only a few eigenmodes with similar taxa clustered into groups; while interactions between taxa within these groups correspond to slow modes [27]. On the one hand, recent experiments show that while the abundances of strains under competition fluctuate and show great compositional variability across samples, the coarse-grained compositions based on functional groups [28][29][30][31] or at family taxonomic levels largely converge (see Fig. 1) [32]. Algorithms have been developed to identify these clustered groups [33][34]. On the other hand, also relevant to this discussion in terms of clustering is the fine-scale structure observed in ribotypes, which refers to unique 16S ribosomal sequences: most ribotypes fall into discrete clusters with 99% sequence consensus [35]. It has been shown that even for closely related strains with 99.99% genome similarity (around 100 base-pairs differences) can have decoupled dynamics observed in abundance time series data due to changes in phenotypes, thus higher composition variability is exhibited at the strain level than at the species level [36]. This opens up spaces for how we think about coarse-graining: perhaps diversity limited by the competitive exclusion principle should not be based on individual strains, but rather on the effective diversity based on clustered groups, (or as we call it here as “clusters” which are composites of multiple “taxa” that not only exhibit similarities but also differences in phenotypes).

**FIG. 1:**
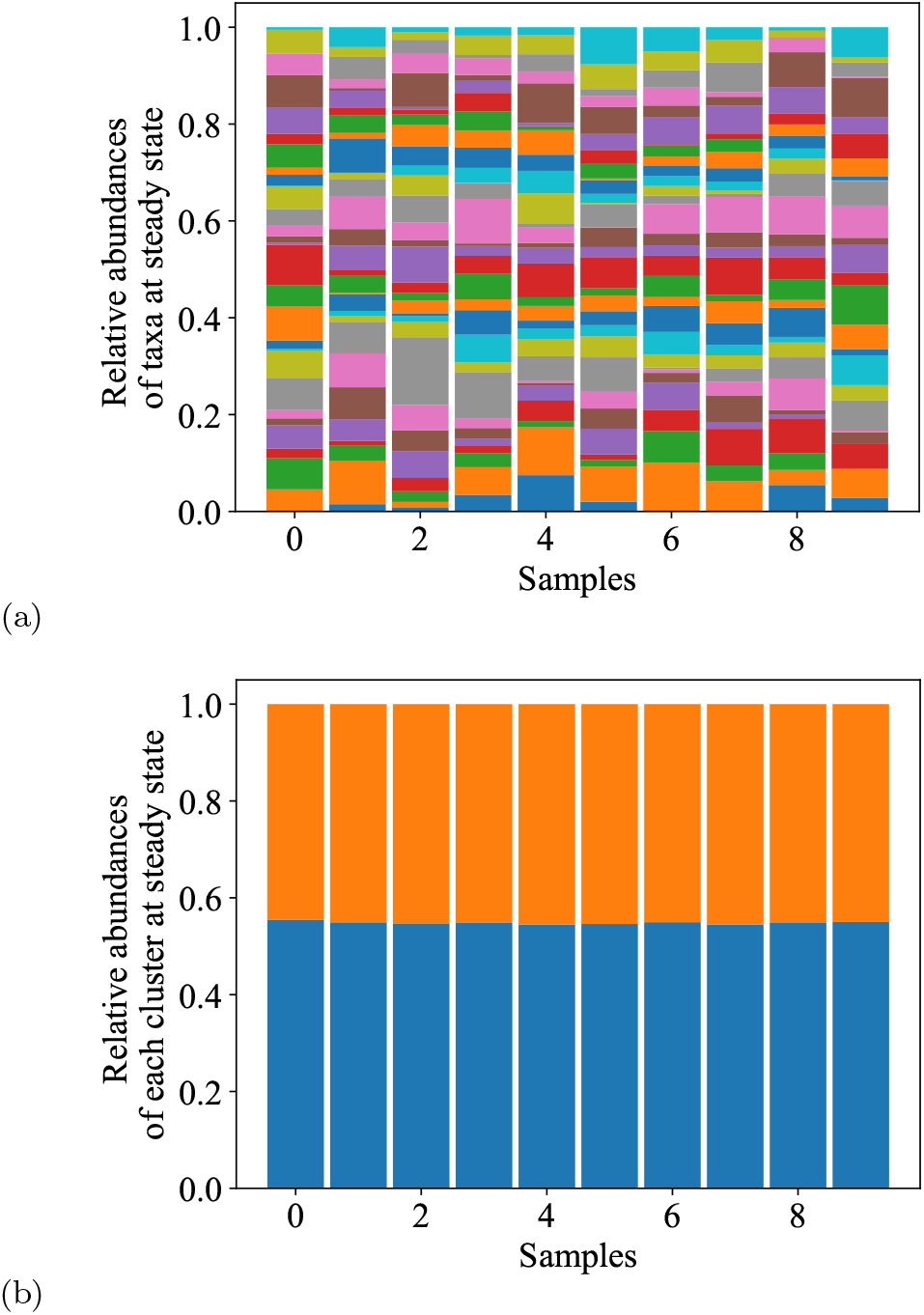
Compositional variability across samples at different taxonomic levels suggests coarse-grainability. As shown in prior experiments (e.g. [31][32]), variability is reduced at a coarse-grained level. (a) Given the abundances *x*_*i*_ of different taxa *i*, the relative abundances *x*_*i*_*/*Σ_*j*_*x*_*j*_ (each taxon notated by a color) vary across samples. (b) On the coarse-grained level, the relative abundance of the two clusters (orange and blue) is consistent across samples. The plots show the taxa abundances at steady state in synthetic data generated by our model depicted in Sect. III B, at different initial conditions.

In this work, we suggest that the shielding effect requires a closer scrutiny and has several general implications outside of prior work. Using the model adapted from [37][38][39], we first present theoretical work to show how the shielding effect is exhibited not only for alternative resources, but also for competitions over essential resources, which has not been previously considered. The model is then extended to more general modes of interactions to consider a mix of both types of resources. We further propose the novel idea that trade-offs, contrary to the previous thinking which consider the upper bound of coexisting species to be lifted [19][20][21][22], offer an alternative way to reassess diversity and reinforce the upper limit: a great number of taxa can be grouped into a small, limited handful of clusters, which in turn satisfy the competitive exclusion principle such that the number of clusters are limited by the number of coarse-grained resources. The dynamics is largely determined on the cluster level through competitive exclusion, while the fitness landscape for the taxa within each cluster is flat, showing a fine-scale structure that supports the coexistence of any arbitrary number of taxa. Using a hierarchy of models, we show in Fig. 2 the coarse-graining through trade-offs at three different levels: (a) all taxa coexist as a single cluster; (b) diversity within each cluster is sustained through shielding effect, while the diversity on the cluster level is limited by the number of coarse-grained resources, thus satisfying the competitive exclusion principle and; (c) multiple clusters self-organize through a second level of shielding, such that there can be more clusters to coexist in the system compared to the number of coarse-grained resources which violates the competitive exclusion principle. Lastly, we show how trade-offs can sustain a community with high diversity that exhibits a hierarchical structure.

**FIG. 2:**
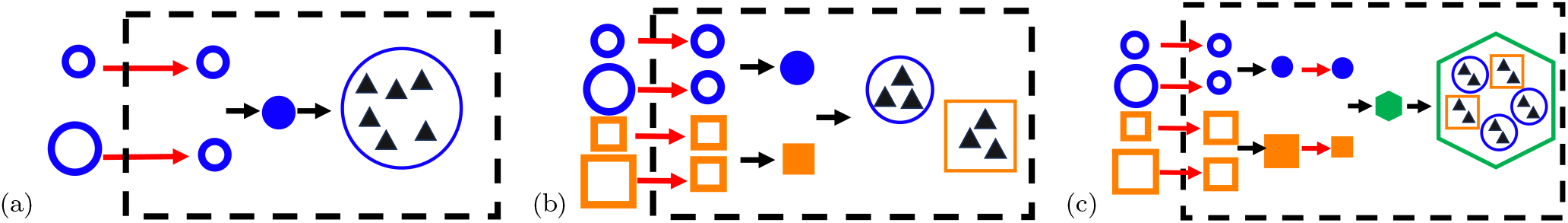
An illustration of three levels of coarse-graining through trade-offs. (a) Previous work [19][20] suggests that given the difference in external supply of resources 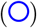 is not too large (with concentrations scale with the size of the markers), the community can self-organize to shield off this asymmetry to bring the resource level in the system (dotted box) to similar levels. We call this shielding effect (red arrow), which can sustain a large system with high diversity. We suggest this can be seen as a small system, where multiple supplied externally resources are effectively coarse-grained into a single growth-limiting factor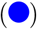, and multiple taxa within the system 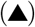 are behaving as one single cluster. (b) By extension, if the externally supplied resources 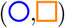 can be coarse-grained into two growth-limiting factors 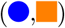, the system can support at most two coexisting clusters, thus satisfying the competitive exclusion principle on a coarse-grained level. This leads to a reduction of compositional variability as shown in Fig. 1. (c) Further coarse-graining of the two growth-limiting factors 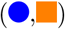 into a single resource 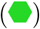 and combines all clusters into a macro-cluster 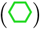. This removes the upper limit of the number of coexisting clusters, and diversity is sustained through a hierarchical structure.

### II. THE MODEL

*A well-mixed system* is considered in this work to study the contributing factors to diversity that are isolated from spatial niche partitioning, which has been well-explored in other work [40][41][42][43][44] and undoubtedly also plays a role in diversification; thus it is not the focus in this paper and we only focus on systems with no spatial heterogeneity. Our system consists of a chemostat, which is a low-cost, automated, continuous culture device that shows no appreciable spatial structures [45][46], and has been used to model a wide range of environments including gut microbiota [47][48] In a chemostat, a fresh medium with concentration *a*_*j*_ of resource *j* is externally supplied continuously while the stale culture liquid is pumped out, both at a flow rate of *F*. The models used in this paper consider bacterial growth in a chemostat that follows the dynamics described by

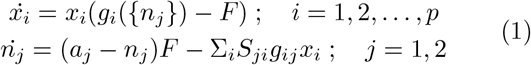

where *x*_*i*_ is the taxon abundance and *S*_*ji*_ is the stoichiometry of converting resources *j* into biomass for taxon *i*. Specifically, we consider a population with *p* taxa that interact only through nutrient consumption, such that the bacterial growth rate *g*_*i*_ depends only on resource levels *n*_*j*_. The growth rate of taxon *i* contributed by the utilization of resource *j* is *g*_*ij*_. In the hierarchy of models used in this paper, we vary the functional forms of growth rates *g*_*i*_ and *g*_*ij*_ for different types of resources involved. We derive these rates in the context of proteome partitioning.

*A coarse-grained model of proteome partitioning* has been proposed to explain the robust empirical laws observed between bacterial growth rates and macromolecular composition, such as protein and RNA content [37][38][39]. Adapting from this line of work, we introduce here a model of bacterial growth that allows us to consider the interactions through competition over alternate resources, essential resources, or a mix of both types of resources. To briefly summarize our model, protein biosynthesis involves nutrient metabolism and protein translation, of which the rates depend on the fraction of the proteome allocated to these processes. For example, the metabolic rate is proportional to the fraction of transporter proteins and enzymes defined as Φ_*M*_, and the translation rate is proportional to the fraction of ribosomal proteins Φ_*R*_ within the proteome. The inset in Fig. 3a shows the proteome constraint highlighted by [38]: while Φ_*R*_ and Φ_*M*_ vary in different environments to optimize bacterial growth, these changes are subjected to the constraint 1 − Φ_*o*_ = Φ_*R*_ + Φ_*M*_, where Φ_*o*_ is fixed and does not depend on the growth rate. Under the proteome constraint, the bacterial growth rate can be derived by assuming flux balance [38][11].

**FIG. 3:**
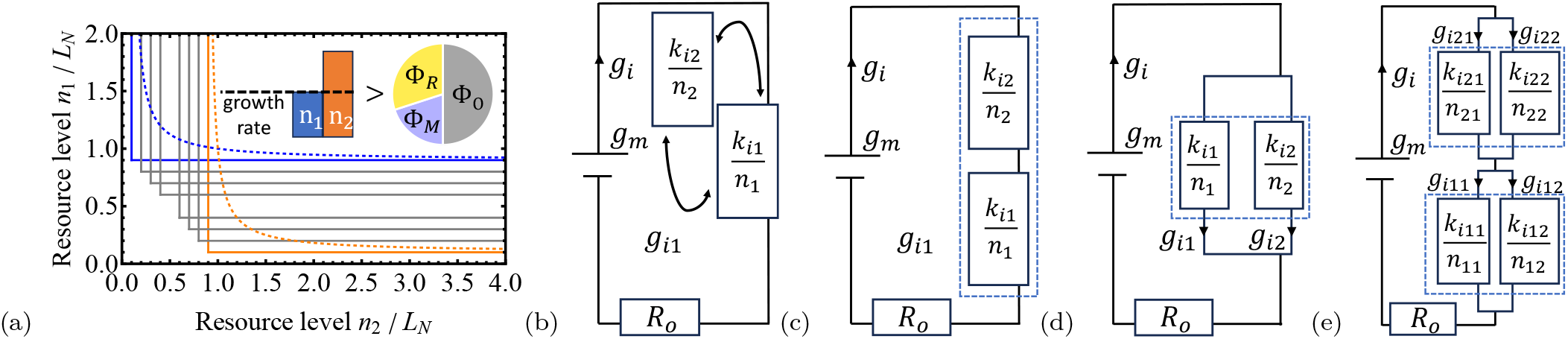
Growth rate based on constraints in proteome: (a) Growth isoclines for different taxa competiting for essential resources under the Liebig’s law (solid lines) and proteome constraint (dotted lines) are plotted. Gray denotes generalists, while blue and orange correspond to the specialists of the two resources (*n*1 and *n*2). Inset refers to the two models. Left: growth rate determined by the most depleted nutrient. Right: proteome fractions Φ_*R*_ and Φ_*M*_ change with the environment, subjected to the constraint that the fraction Φ_*O*_ is fixed. This results in a slower growth rate. The circuitry analogy for (b) the Liebig’s law, and for proteome constraint under (c) interactive essential resources, (d) alternate resources, and (e) a mix of both. The resistance *k*_*ij*_*/n*_*j*_ increases when resource *j* is depleted, leading to a small growth rate *g*_*i*_. Trade-offs in (10)-(11),(17) are notated by dotted boxes.

The functional form of the growth rate derived from proteome partitioning has the same form as Ohm’s law in circuitry, as shown in [37][39]: the growth rate is affected by the growth-limiting resources in a way similar to how the current in a circuit is reduced due to resistors. In this analogy, the current is the growth rate *g*_*i*_ at the exponential phase. The voltage is the maximum growth rate *g*_*m*_, which is proportional to the total available resources Φ_*R*_ + Φ_*M*_. The total sum of resistance is *R*_*o*_ +Σ_*j*_*R*_*ij*_, where *R*_*o*_ is fixed and independent of the nutrient levels (e.g. rate of translation is restricted by the amount of ribosomes) and Σ_*j*_*R*_*ij*_ describes the effects of the growth-limiting factor *j* for strain *i*. Similar to the Ohm’s law (current = voltage */* resistance), the growth rate of taxon *i*, as shown in SM Sect. 1, takes the form of

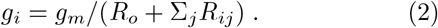

This analogy of using electric circuits to describe the bacterial growth rate is shown in Fig. 3c-e. In hindsight, this is not surprising as the proteome model describes the allocation of resources to different processes; whereas in a closed circuit, the total sum of voltage dropped across each resistor is conserved and is provided by the electromotive force of the battery. This coarse-grained model allows us to adopt a modular approach of combining different loads, where the drop in the resource level increases the resistance (*R*_*ij*_ = *k*_*ij*_*/n*_*j*_), where 1*/k*_*ij*_ is proportional to the efficiency of taxon *i* at growing on nutrient *j* [11], and its biological interpretation is explored in SM Sect. 1. Depending on the circuitry, we can now consider the cases of interactive essential resources, alternate resources, and a combination of both types of resources.

*Essential resources* refer to a set of resources that are all required for bacterial growth. They can be further classified into two categories: non-interactive and interactive [49]. The prior follows the Liebig’s law of minimum [50], which states that the growth rate is limited only by the scarcest of the resources. As shown in Fig. 3a, the growth rate isoclines for the taxa (*g*_*i*_ = constant) are non-differentiable functions of the resource levels. This is akin to the circuitry in Fig. 3b of having a single resistor that is swapped in when that corresponding resource becomes scarce and growth-limiting. In contrast, interactive essential resources result in the growth rate isoclines in Fig. 3a that are differentiable; they asymptotically approach the non-interactive case but differ at intermediate levels of the resources. Previous work related to tradeoffs, such as [22], considers essential resources only for the non-interactive case, and therefore did not consider the shielding effect in the context of essential resources.

Under the proteome constraint, we show in SM Sect. 1.2 that the growth rate functional in Eq. (2) with two essential resources takes the following form:

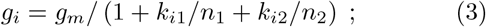

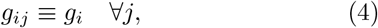

where *n*_1_ and *n*_2_ are the levels of the two resources with scaling constants *k*_*i*1_ and *k*_*i*2_, and *g*_*ij*_ is the bacterial growth rate due to resource *j* for taxon *i*, which is the same across resources since growth is limited by all resources. This is analogous to the circuitry in which the resistors are connected in series with *R*_*o*_ = 1, as shown in Fig. 3c: the model corresponds to the interactive case due to changes in proteome partitioning as the environment changes, and the resulting total resistance is higher than that in the non-interacting case, leading to a smaller growth rate. In the next section, it will become apparent that this drop in growth rates at intermediate levels of nutrients is important to maintaining diversity.

*Alternate resources* refer to a set of resources of which at least one has to be present to sustain bacterial growth. For the system involving two alternate resources, we show in SM Sect. 1.1 that the functional form of the growth rate in Eq. (2) can be written as

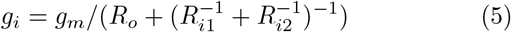

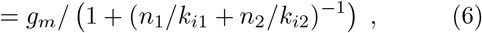

This is analogous to the case of resistors connected in parallel in Fig. 3d. The total growth rate *g*_*i*_ is contributed by *g*_*ij*_ from different resources *n*_*j*_, such that both the Kirchhoff’s current law *g*_*i*_ = Σ_*j*_*g*_*ij*_ and the relation of *g*_*i*1_*R*_*i*1_ = *g*_*i*2_*R*_*i*2_ should be satisfied, as in a parallel circuit. This can be achieved if the rates *g*_*ij*_ take the form of

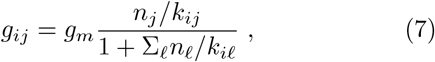

with *j*, ℓ = 1, 2 for a model with two resources.

*Mixed modes of interaction* consider competitions involving both alternate and essential resources. Eq. (1-7) consider specific cases that consist of only one of the two modes. But for the general case with a mixed of modes, as shown in Fig. 3e, the growth rate of taxon *i* contributed by essential resource *e* = 1, 2, can be written in terms of resistances *R*_*ie*_

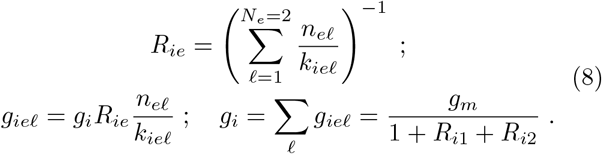

For essential nutrient *e*, there are multiple alternate sources each with concentration *n*_*eℓ*_, with *ℓ* = 1, …, *N*_*e*_.

**TABLE I:** Model Parameters.

| Parameters | Notations |
| --- | --- |
| Growth-related proteome fraction | $1 - \Phi_o$ |
| Metabolic proteome fraction | $\Phi_M$ |
| Ribosomal proteome fraction | $\Phi_R$ |
| Bacterial growth rate of taxon $i$ | $g_i$ |
| Maximum growth rate in optimal environment | $g_m/R_o$ |
| Level of resource $j$ (alternate / essential) | $n_j$ |
| Michaelis-Menten constant for level of resource $j$ | $k_{ij}$ |
| Resistance due to limitation of resource $j$ | $R_{ij} = k_{ij}/n_j$ |
| Level of alternative source $\ell$ for resource $e$ | $n_{e\ell}$ |
| Growth rate due to alternate source $\ell$ of resource $e$ | $g_{iel}$ |
| Michaelis-Menten constant for alternative source $\ell$ of resource $e$ | $k_{iel}$ |
| Number of alternate sources for resource $e$ | $N_e$ |
| Stoichiometry between resource $j$ and taxon $i$ | $S_{ji}$ |
| Constants in trade-offs | $\tilde{k}, \tilde{k}_{Ie}$ |
| Flow rate of the chemostat | $F$ |
| Supply rates of resource $j$ | $a_j$ |
| Impact factor | $m_{T,N}^{(i)}$ |
| Scaling for alternate resources | $L_N = [(g_m/F - 1)\tilde{k}]^{-1}$ |
| Scaling for essential resources | $L_N = (g_m/F - 1)^{-1}\tilde{k}$ |
| Impact ratio | $m_{1,2}^{(i)}$ |
| Interaction matrix | $b_{ij}$ |
| $k_{ij}$ for specialist 1 | $k_{i'j}$ with $i' = \operatorname{argmin}_i k_{i1}$ |
| $k_{ij}$ for specialist 2 | $k_{i'j}$ with $i' = \operatorname{argmax}_i k_{i1}$ |
| Average self-interaction strength | $b_s = (b_{11} + b_{22})/2$ |
| Average cross-interaction strength | $b_c = (b_{12} + b_{21})/2$ |
| Symmetry breaking terms | $s = (b_{22} - b_{11})/2$ |
| | $e = (b_{12} - b_{21})/2$ |
| Stabilization factor | $b_s - b_c$ |
| Equalization factor | $e - s$ |
| Test statistic for augmented Dickey-Fuller test | $\tau$ |
| Abundance of taxon $i$ | $x_i$ |
| The set of taxa under cluster I | $H_I$ |
| The set of all taxa in the system | $H_o$ |
| Abundance of cluster I | $X_I = \sum_{i \in H_I} x_i$ |

The last two relations ensure that Kirchhoff’s current law is satisfied. To reduce the number of parameters, we assume that all alternate sources *ℓ* of each essential resource *e* share the same stoichiometry *S*_*ei*_, such that Eq. (1) is modified as

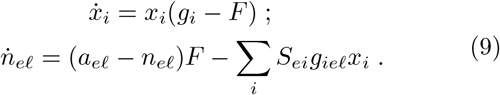

## III. RESULTS

### A. First-level of coarse-graining: the shielding effect is applicable not only to alternative resources but also to interactive essential resources

Using models of competitions over only alternate or essential resources (Figure 3c-d), we consider in this section the first level of coarse-graining with one single cluster composed of multiple taxa, as illustrated in Fig.2a.

#### Trade-offs allow equalization of growth rates by re-balancing of resource levels

In competitions over alternate resources, trade-offs can lead to rich diversity sustained by the shielding effect, as shown in previous theoretical work [19][22][51]. Given that the growth rate functionals in our model are different, we first demonstrate that previous results on alternate resources are also applicable to our model, as defined by Eq. (1) and Eq. (6-7). Fig. 4a shows the zero net growth isoclines, or ZNGIs defined as *g*_*i*_(*n*_1_, *n*_2_) = *F* for each of the competing taxa over two alternate resources in a chemostat: they define a set of environmental conditions (*n*_1_, *n*_2_) under which the taxa can survive at steady state. We notice that if the following condition holds:

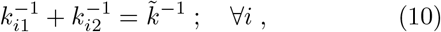

then the ZNGIs intersect at the scaled resource levels of 1 *≡ n*^*⋆*^*/L*_*N*_ = *n*_1_*/L*_*N*_ = *n*_2_*/L*_*N*_, with no fine-tunings of parameter required. The length scale for the alternate resources *L*_*N*_ is defined by Eq. (11) in SM Sect. 2. The condition in Eq. (10) can be understood as a trade-off between the competitiveness for the two alternate resources: a taxon that is more competitive for resource 1 would correspond to a larger value of 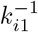, and therefore a smaller value of 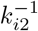 which is associated with a lower competitiveness for resource 2. This is interpreted as the constraint of resource allocation within Φ_*M*_ for the two resources, as discussed in SM Sect. 1.1.

**FIG. 4:**
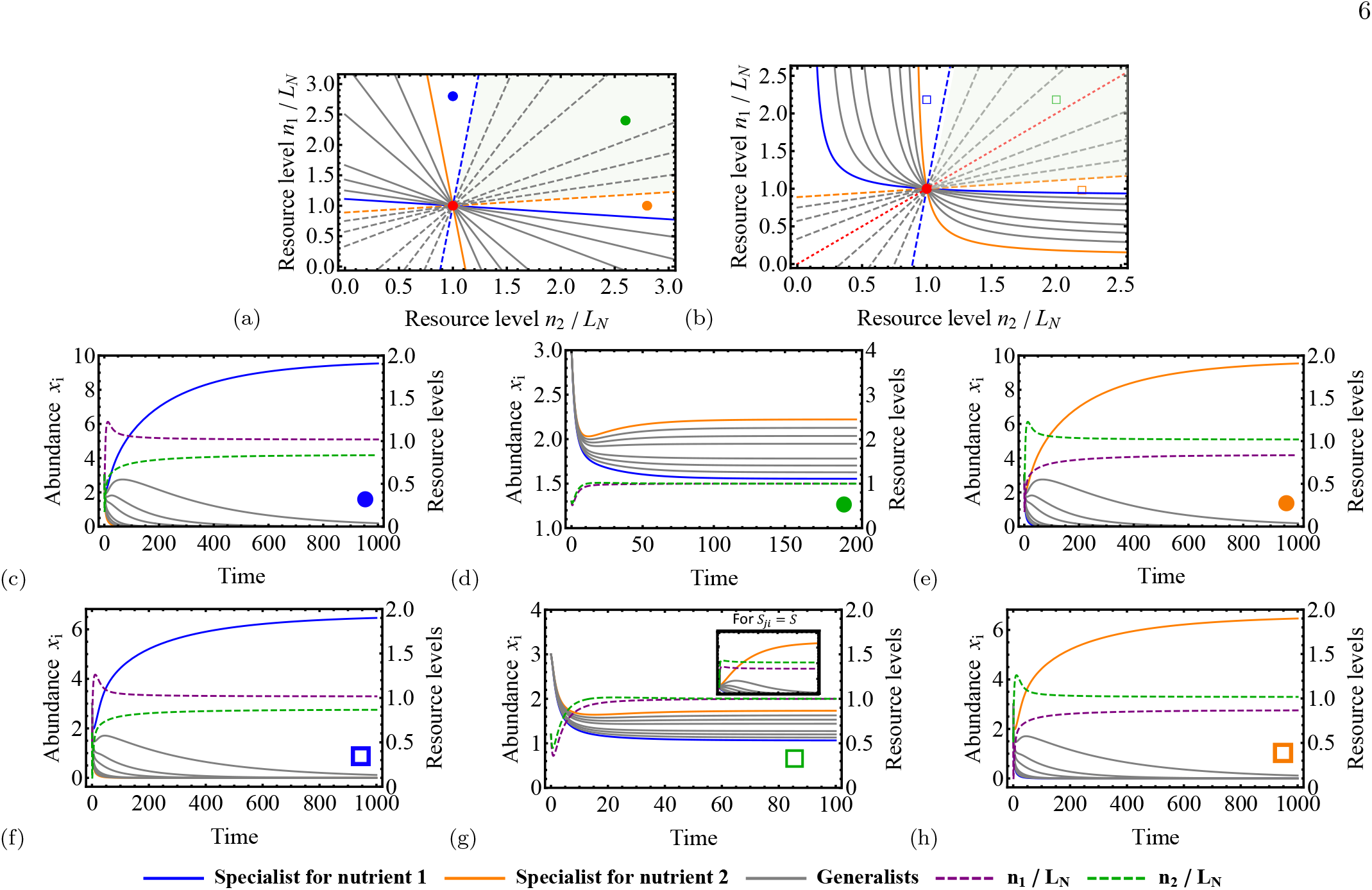
Comparison between competitions for alternate resources (a, c-e) and interactive essential resources (b, f-h) with different supply concentrations. (a-b) Zero Net Growth Isoclines (ZNGIs) (solid) and impact lines (dashed) for the specialists (blue, orange) and generalists (gray) are plotted in the resource space. Dashed red line shows impact lines for all taxa when there are no trade-offs in stoichiometry with nutrient competitiveness (*S*_*ji*_ = *S*). Depending on whether the supply concentrations (*a*_1_, *a*_2_), denoted by the colored markers 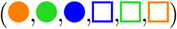, fall within the convex hull (green shaded region in a-b), the competition outcome is either coexistence or competitive exclusion. For *S*_*ji*_ = *S*, the convex hull has zero area, leading to competitive exclusion in the subset of fig.4g.

The solution of *n*_*j*_*/L*_*N*_ = 1 to solving *g*_*i*_ = *F* for all taxa corresponds to the environmental condition at which all taxa would share the same growth rate at 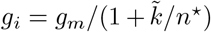. The taxa abundances and resources levels over time are obtained through numerical integration and are plotted in Fig. 4d, which shows that all taxa indeed coexist at steady state, with the scaled levels for both resources approaching unity. In other words, despite the differences in resource external supply rates (*a*_1_*/L*_*N*_ *a*_2_*/L*_*N*_), the microbial community can self-organize to shield the internal environment from the external asymmetry to re-balance both resources to the same levels (*n*_1_*/L*_*N*_ = *n*_2_*/L*_*N*_). We call this the shielding effect. In this environmental condition, both resources are treated as equals and no fitness advantage can be gained by utilizing a specific strategy defined by the values of (*k*_*i*1_, *k*_*i*2_).

Given that trade-offs in competitiveness over essential resources, such as carbon and phosphate, have been observed in nature [52][53], do trade-offs for essential resources also lead to equalized bacterial growth rates at shielding? For a system where taxa compete for two essential resources as defined by Eq. (1) and Eq. (3-4), the ZNGIs intersect if the following condition is satisfied:

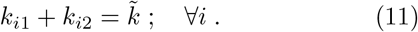

Eq. (11) describes a trade-off relation: for a taxon that is highly competitive for nutrient 1 (with a small *k*_*i*1_), its competitiveness for nutrient 2 is low (with a large *k*_*i*2_), as discussed in SM Sect. 1.2. The ZNGIs for the case of essential resources are plotted in Fig. 4b. Unlike the case of alternate resources, the ZNGIs are non-linear. Nonetheless, the interactiveness due to proteome partitioning allows the ZNGIs to intersect at *n*_1_*/L*_*N*_ = *n*_2_*/L*_*N*_ = 1, where the length scale for essential resources is *L*_*N*_ as defined by Eq. (S15) in SM Sect. 2. This re-balancing of the resources to the same level through shielding is not possible for the non-interactive case (see Fig. 3a). Consequently, taxa with different preferences for either of the resources are equally optimal. The time series obtained from numerical integration in Fig. 4g show that, given shielding, high diversity can be sustained at steady state by only two essential resources. This is in contradiction to the competitive exclusion principle which states that at most only two taxa can coexist.

#### Only small asymmetry in resource supplies can be shielded

Although shielding (or the re-balancing of resource levels) is possible for both modes of resource competition, it does not necessarily mean that the system will always reach shielding — the community can still collapse. It depends on whether the bacterial community as a consortium can shield the resource levels (*n*_1_, *n*_2_) from the asymmetry in the resource supplies (*a*_1_*/L*_*N*_ ≠ *a*_2_*/L*_*N*_), of which the mechanism can be understood in terms of resource consumption. From Eq. (1), we define for taxon *i* the impact factor as the ratio of consuming resource 1 to resource 2:

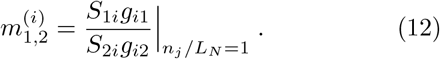

Then for each taxon, we define the impact line as 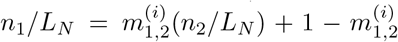, which are plotted in Fig. 4b-B. The impact line is a straight line passing through the steady-state solution (*n*^*⋆*^*/L*_*N*_, *n*^*⋆*^*/L*_*N*_) with slope 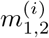. The high diversity state can be reached only when the supply vector 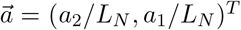 falls within the shaded region bounded by the impact lines, as shown in Fig. 4d,g. If asymmetry in source supply is too large and the supply vector falls outside of the region, the fittest taxon would out-compete all other weaker taxa and dominate the population, resulting in competitive exclusion. For both alternate resources (Fig. 4c,e) and essential resources (Fig. 4f,h), the collapse of diversity due to the failure to reach shielding is accompanied by the separation of resource levels (*n*_1_*/L*_*N*_ ≠ *n*_2_*/L*_*N*_). We show the derivation of this requirement for coexistence in SM Sect. 2. In the context of alternate resources, this bounded region is equivalent to “the convex hull” [19]: the resource supply concentrations are bounded by the Michaelis-Menten constants, or *a*_*j*_ ≤ max(1*/k*_*ij*_) for all resources *j*. In other words, the system requires specialists (with extreme values of *k*_*ij*_) to bring the system to shielding. For essential resources, we will show in the next section that this statement of convex-hull applies when there is an additional implicit trade-off.

#### An implicit trade-off between nutrient efficiency and impact ratio stabilizes coexistence

In the context of alternate resources, an implicit assumption taken in past work [20][22][51][49][54] is that there exists another implicit trade-off such that taxa which are more adapted at growing on a particular resource would also consume more of that resource. This is true when the impact factor satisfy the following relation:

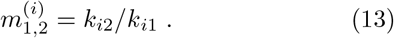

Under this implicit relation, a taxon that is highly competitive for resource 1 (with a small *k*_*i*1_) would have a large impact factor 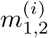, which indicates that this taxon consumes more of nutrient 1. Geometrically, Eq. (13) means that the impact lines are perpendicular to the ZNGIs and specialists of resource 1 would correspond to an impact line with a steeper slope, as shown in Fig. 3A. Based on Eq. (7) and Eq. (12), the implicit trade-off Eq. (13) can be incorporated into the model by setting the stoichiometry over all resources to be the same for all taxon (i.e. *S*_*ji*_ = *S* ∀_*i*,*j*_), as seen in previous models [20][22][51].

To show diversity can be developed through shielding in systems with essential resources, we apply the same assumption of Eq. (13). From Eq. (4) and Eq. (12), this means the stoichiometries and the nutrient efficiencies follow the relation: *S*_*ji*_ = 1*/k*_*ij*_. This relation is plausible in a sense that in theory, it is clear that coexistence in a competition with two taxa requires the taxon that grows better at a particular resource (with a smaller *k*_*ij*_) to have a larger stoichiometry of that resource [49][54]; experimental work further suggests a trade-off relation between efficiency and yield [55]. This implicit assumption of the relation between nutrient efficiency and impact ratio in the form of Eq. (13) sets up a “convex hull” for both alternative and essential resources by the specialists as shown in Fig. 3a-b, and stabilizes the shielding effect. Both diagrams are topologically equivalent locally in the vicinity of the coexistence solution: a taxon with a smaller slope of ZNGI has a steeper slope of the impact line. Notice that we have not introduced new assumptions to the model of essential resource that are not present in models of alternative resources in previous work [19][22][51]: both classes of models require making assumptions on the stoichiometries *S*_*ji*_ such that the trade-off in Eq. (13) holds.

The stability of the system with essential resources is sustained by the convex hull due to this implicit trade-off in Eq. (13). In the inset of Fig. 4g, we show the time series under a different assumption on the stoichiometries: *S*_*ji*_ = *S*_*j*_ for all *i* such that Eq. (13) no longer holds. The immediate consequence is the collapse of the community. The reason is that the convex hull in this case has no volume. For essential resources, combining Eq. (3) and Eq. (12) gives an impact ratio of 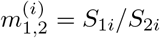. Assuming *S*_*ji*_ = *S* as we did with the alternate resources, the resulting impact ratio is *m*_*i*_ = 1, which is independent of the strategies (*k*_*i*1_, *k*_*i*2_) adopted by the taxa. In other words, every taxon would have the same impact factor, the ZNGIs for all taxa overlap and the bounded region has a zero area, as shown in Fig. 4b. Under this case, coexistence can be sustained only if the resource supplies (*a*_1_, *a*_2_) are fine-tuned to match exactly with the impact lines. Perhaps this is the reason why the shielding effect has never been considered for essential resources until now.

#### Fate of community is determined by community structure in the Lotka-Volterra framework, and points to a lower-dimensional description

While having access to the full model described in Sect. II would allow good predictions on competition outcome, not all growth-limiting factors, such as all resource levels, are traced in real data. Nonetheless, effective couplings between taxa can still be inferred from the time series of taxa abundances [56][57][58][59]. Naively, we can assume that the dynamics of the taxa abundances *x*_*i*_ follows the competitive Lotka-Volterra Equations

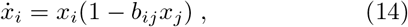

which can be understood as a local expansion of the full system up to second-order terms. The interaction matrix *B* consists of positive constants *b*_*ij*_ *>* 0, which describe effective couplings between taxa, mediated through resource levels as described in the full model.

To capture the inter-taxon interactions, we approximate the full system described by Eq. (1) for both essential and alternate resources using competitive Lotka-Volterra Equations, as derived in SM Sect. 3. To briefly summarize: we take the adiabatic approximation 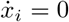 in Eq. (1) to solve for the resource levels *n*_*j*_, which are then substituted in the set of equations 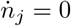, giving the couplings between taxa. For essential resources, the community matrix depends on the resource supplies (*a*_1_, *a*_2_), and has the following expression:

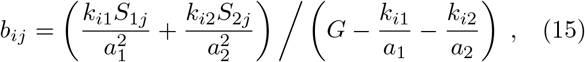

where *G ≡ g*_*m*_*/F* − 1.

Does the Lotka-Volterra approximation show the same transitions in diversity in Fig. 4? And how do these transitions relate to the community structure? Using Eq. (15) to approximate the dynamics of the three cases in Fig. 4f-h with different resource supplies (*a*_1_, *a*_2_), the taxa abundances are plotted in Fig. 5 for the case of essential resources. A similar plot is made for alternate resources in Fig. S2 of SM Sect. 3.1. The predictions of the competition outcomes from these approximations qualitatively match with the full system. Outside of shielding, competitive exclusion drives down the diversity such that only one taxon dominates. The winning taxon, which is always a specialist, corresponds to the row in the interaction matrix where the coupling terms are the weakest, as shown in the insets of Fig. 5a, c. By comparison, the losing taxa are suppressed more strongly in the presence of other taxa. Interestingly for the high diversity state in Fig. 5b, where both specialists and generalists coexist, sub-community structures emerge. The population can be separated into two clusters as shown by the block matrix in the subset. This begs the question: is there a description of this system simpler than a model that requires the number of parameters to scale with *O*(*N*×*N*)?

**FIG. 5:**
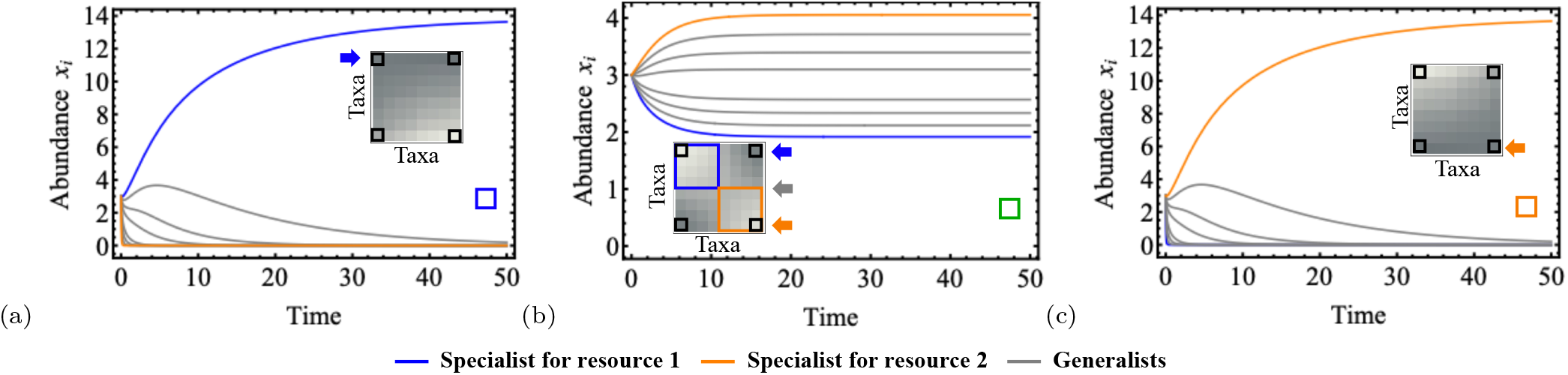
Taxa abundance time series from the Lotka-Volterra approximation under competitions for essential resources, with three different resource supplies that correspond to Fig. 4b,f-h, showing qualitative agreements with full system. Inset shows the interaction matrix *B*, where a lighter value means a stronger coupling *B*_*ij*_, and the arrow shows the rows for the winning strain in the case of competitive exclusion. Using the matrix elements at the corners (in black square), we model this in Fig.6 as a competition between the two specialists (blue and orange) while ignoring the generalists (gray), with four possible outcomes.

#### Redundancy in generalists: dynamics of the entire community can be described only by interactions between specialists

It turns out that for a competition over two resources, the outcome does not depend on the generalists and we can well predict the diversity of the full system using a Lotka-Volterra framework with a much smaller model that involves only two taxa: specialists of the two resources correspond to argmin_*i*_*′ k*_*i*_*′*_1_ and argmax_*i*_*′ k*_*i*_*′*_1_. Specifically, we consider a new 2 × 2 interaction matrix *b*_*ij*_ for the two specialists as defined by Eq. (15), meaning that this smaller interaction matrix now consists of matrix elements notated by the black squares in Fig. 5a-c.

We now look at the competition between the two specialists (Fig. 6a). Interaction matrix *B* can be reparameterized in terms of the average self-interaction strength *b*_*s*_, the cross-interaction strength *b*_*c*_, and the symmetry-breaking terms *s* and *e* such that

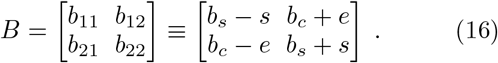

**FIG. 6:**
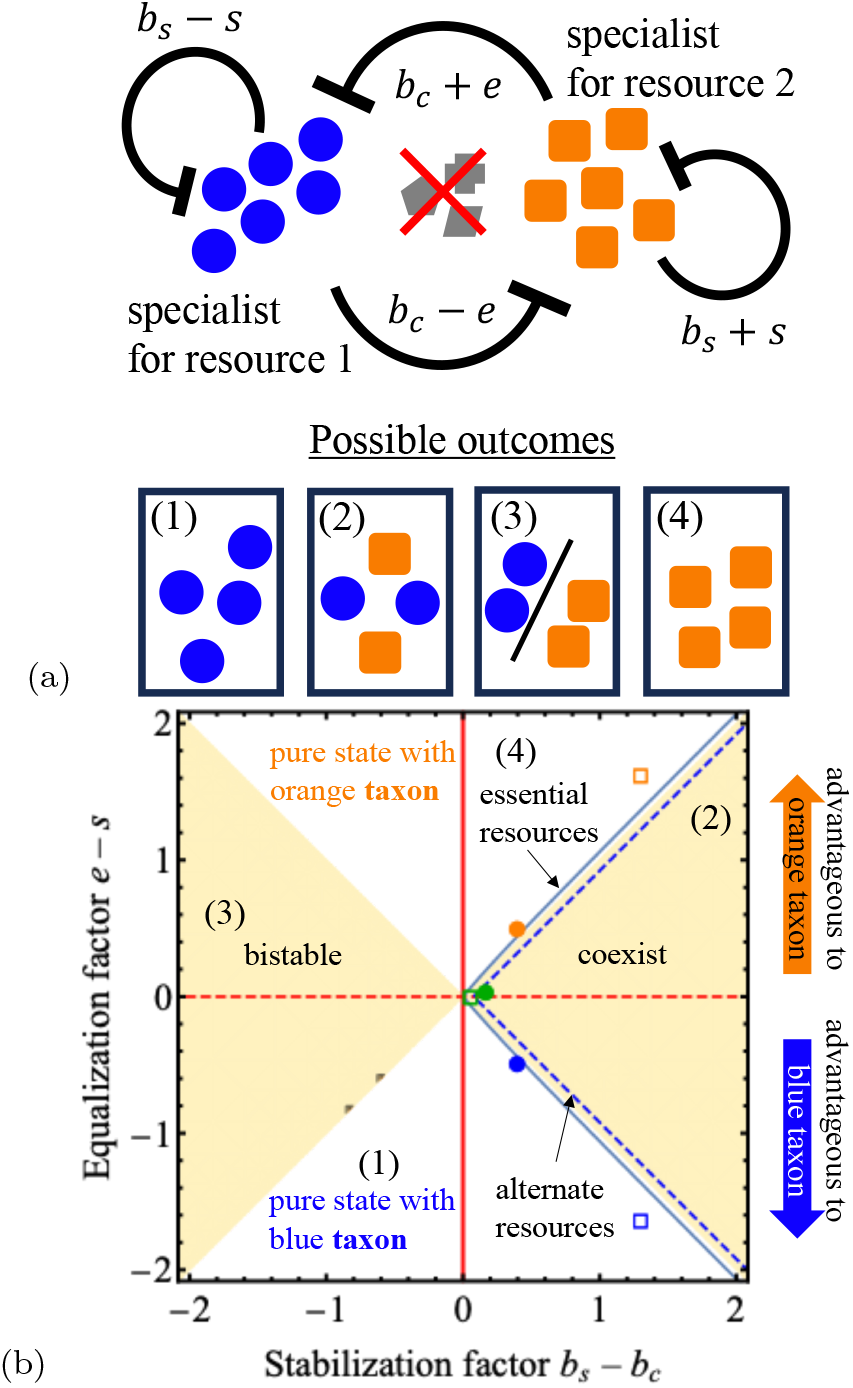
Competition between two specialists can explain transition of diversity in the full system under the first level of coarse-graining (Fig. 2a). (a) A two-taxon description under Lotka-Volterra approximation. (b) The corresponding phase diagram can be partitioned into four regions. The solid and dotted blue lines show the boundaries for transitions of diversity derived from the full model with multiple taxa. The scatter plot corresponds the six cases in Fig. 4. Only the shielding cases with high diversity (green) fall inside region (2).

In general, Lotka-Volterra models considering competitions with two taxa have been very well-studied. Our scheme of re-parametization allows us to understand the system in a framework related to [60][61], but now that the phase diagram in Fig. 6b is instead partitioned by linear boundaries into four sub-regions that correspond to different outcomes. Specifically, the system is determined by two parameters: the equalization factor |*e* − *s*| and the stabilization factor *b*_*s*_ − *b*_*c*_. There are four possible outcomes : (1,4) competitive exclusion by one of the specialists; (2) coexistence and; (3) bistability where one of the specialists is driven to extinction, depending on initial conditions. As derived in SM Sect. 3.3, these linear boundaries are given by |*e*− *s*| = |*b*_*s*_ − *b*_*c*_| : coexistence is supported only when the magnitude of the equalization factor |*e* − *s*|, which describes the asymmetry of the two taxa, is small. As the magnitude of the equalization factor increases, coexistence can be sustained only when the stabilization factor *b*_*s*_ − *b*_*c*_ is large. In other words, given a large asymmetry between the two taxa as characterized by the equalization factor, they can still coexist provided that self-interaction is strong enough to stabilize the system compared to the destabilizing force due to inter-taxon competition.

Given the varying resource supply concentrations (*a*_1_, *a*_2_) in Fig. 4, we use (15) to obtain the two-taxon interaction matrix *B* and identify the corresponding locations in the phase diagram in Fig. 6b. Interestingly, the prediction by this simpler model agrees with the competition outcome in the full model, and both descriptions largely agree on the conditions for coexistence. To obtain the boundaries of transitions in the full model for comparison, we consider the set of supply concentrations (*a*_1_, *a*_2_) that falls on the two impact lines (one for each of the specialists), which defines the convex hull in Fig. 4a-b: the resource supply concentration follows the relation 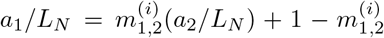. We can then map this set of supply vectors (*a*_1_, *a*_2_) to the stabilization factor *b*_*s*_ − *b*_*c*_ and equalization factors *e*− *s*, which gives the boundaries of the transitions in the two-dimensional phase space as denoted by the blue lines in Fig. 6b. The plot shows slight deviations with the boundaries from the Lotka-Volterra approximation phase diagrams but otherwise largely in agreement. In particular, using Eq. (S24) and Eq. (S27) in SM Sect. 3, we placed on the phase diagram the six cases of Fig. 4 for both alternate resources (circles) and essential resources (squares), with varying resource supplies (*a*_1_, *a*_2_). When the asymmetry in resource supply is small (green), the small differences in the equalization factor *e*− *s* can be overcome by stabilization due to the two specialists. This results in a high diversity. On the other hand, having a large asymmetry in the resource supplies (*a*_1_, *a*_2_), a small flow rate *F*, or a small total supply of resources *a*_1_ + *a*_2_ not only increases the stabilization factor, but also the equalization factor, thus driving the system towards pure states where only a single taxon survives (orange and blue). This helps explain the community structure in the inset of Fig. 5b: while each taxon strongly competes with other taxa within the same cluster, they are weakly coupled to taxa in a different cluster (denoted by the blue and orange squares). This strong “self-interaction” with taxa favoring the same resource stabilizes a highly diverse community, and overcomes difference in the equalization factor between the taxa. This stabilization-against-equalization dichotomy will be further discussed for the second level of coarse-graining in the next section.

### B. Second level of coarse-graining: shielding effect in a competition with both essential and alternative resources

As we have shown, the shielding effect lifts the upper limit of taxa that the environment can support: there can be however many taxa to coexist at steady state. The total biomass, however, is constrained due to the conservation of mass: all cell growth is accounted for by the amount of resources pumped into the system. Another way of seeing this is that all coexisting taxa within the system can be effectively coarse-grained as a single cluster with an abundance determined by the amount resources supplied, as demonstrated in Sect. III A and SM Sect. 4. In other words, shielding provides a linkage between large systems with high diversity and the smallest possible system with only one dominating cluster - this is the first level of coarse-graining as depicted in Fig. 2a. Given that recent work suggests diversity can be coarse-grained based on similarities in phenotypes and functional groups [27][32], can our model support a community structure with more than one cluster?

In this section, we explore how the shielding effect provides a path to developing fine-scale diversity: a community consists of multiple coexisting clusters, each composed of a group of taxa that exhibit very different phenotypes but share the same set of trade-offs. This is the second level of coarse-graining as illustrated in Fig. 2b. Specifically, we considered a system defined in Eq. (8)-Eq. (9), with a total of four growth-limiting resources: two essential resources *e*, each has two alternative sources *ℓ* with concentrations *n*_*eℓ*_, with *e*, *ℓ* = 1, 2. Fig. 3e is the graphical representation of the system.

### The reduction in effective dimension of resources allows the survival of more than one cluster

As an extension from the trade-off over alternate resources in Eq. (10) of Sect. III A, we introduce sub-index *I* to denote the specific trade-offs imposed on cluster *I*, which consists of a group of taxa in the set *H*_*I*_ :

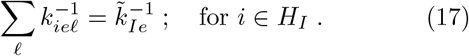

The system we study in this section consists of two clusters *I* = 1, 2, and therefore Eq. (17) corresponds to four constraints defined by 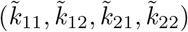. For completeness, we also define *I* = 0 with *H*_0_ = {*i* = 1, 2, 3, …, *M}*, which is the set that includes all taxa such that *H*_*I>*0_ *⊆ H*_0_. The interpretation of Eq. (17) is that various taxa classified under the same cluseter *I* would have the same competitiveness for essential resource *e*, but different preferences towards the alternate sources *ℓ* of that said resource. Similar to the Sect. III A, we assume that taxa within the same cluster share the same stoichiometries which has the following relation with with their competitiveness over essential resources: *S*_*ei*_ = 1*/k*_*Ie*_ for all *i ∈ H*_*I*_.

Suppose the asymmetry in resource supply across alternate resources is small such that the shielding effect is sustained, how does the competition between two clusters reflect on the environment? Using Eq. (8)-Eq. (9) with various resource supplies *a*_*eℓ*_, we plotted the time-series of both the taxa abundances and resource levels in Fig. 7a-c. The immediate consequence of this new set of trade-offs is that at shielding, the resource levels across the alternate channels of an essential resource *e* are the same 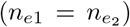. The implication is that, effectively, there are only two resources (*e* = 1, 2) at steady state.

**FIG. 7:**
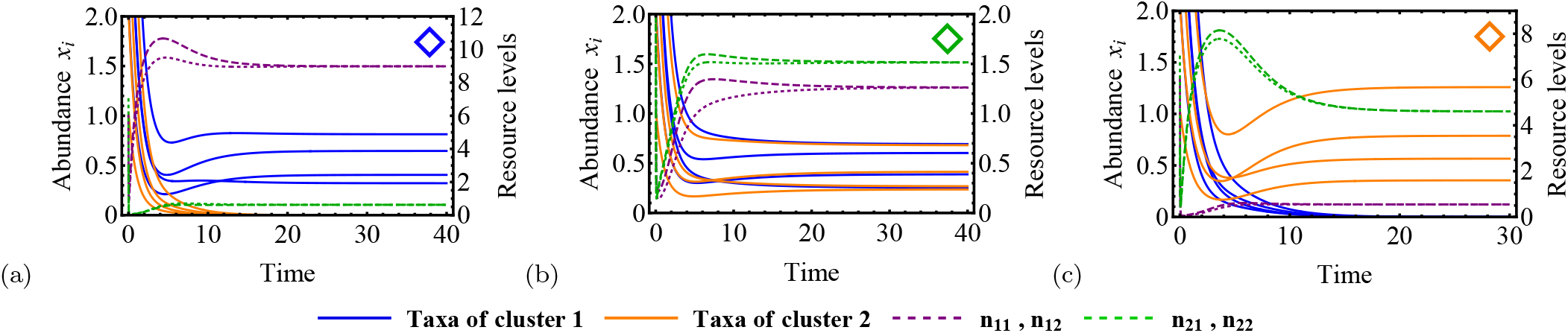
The second level of coarse-graining (Fig. 2B) from taxa to clusters. Abundance involving two different clusters (blue and orange), each with four taxa, competing for four resources (*n*_11_, *n*_12_, *n*_21_, *n*_22_) as supply resource (*a*_11_ = *a*_12_ = *a*_1_*/*2,) changes: (a)(*a*_1_ = 10, *a*_2_ = 30); (b)(*a*_1_ = 18, *a*_2_ = 22) and; (c)(*a*_1_ = 30, *a*_2_ = 10). Dynamics is determined on the cluster level: either coexistence of two clusters, or pure state dominated by one of the two clusters.

The consequence of the trade-offs is that the dynamics exhibits a reduction in dimensionality As supply resources *a*_*eℓ*_ changes, there is a transition from a pure state where all taxa within cluster 1 (blue) coexist and dominate over cluster 2 (orange) in Fig. 7a, to coexistence of both clusters in Fig. 7b, and finally to the extinction of cluster 1 in Fig. 7c. These transitions of competition outcomes occur on the level of clusters, and the taxa that are grouped into the same cluster share the same fate. This is drastically different from the previous section, where it is either competitive exclusion with only one taxon surviving or coexistence of all taxa in Fig. 4. In SM Sect. 4, we show mathematically that the model on the scale of taxa can be rewritten into a model on the scale of clusters. The intuitive picture is that since we have two effective resources, we can have at most two coexisting clusters as stated by the competitive exclusion principle and that the system is essentially a two-cluster competition.

#### Large systems can be explained by effective coupling between clusters in small systems

Similar to the first layer of coarse-graining with only one cluster, we can approximate the full model with a 2 × 2 interaction matrix *B* in Eq. (16). Differing from the prior case in Fig. 4d that depends on two specific taxa which are specialists (Fig. 6a), however, the latter case does not ignore the generalists: the matrix depends on the parameters that define the trade-off and stoichiometries of the clusters in Eq. (17) and describes the competition between all taxa, including both specialists and generalists (Fig. 8a): this interaction between the two clusters *I, I*′ = 1, 2 can be described by the Lotka-Volterra approximation in Eq. (15) with matrix elements of 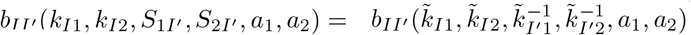. In Fig. 8b, we plotted on the phase diagram of this simplified system the three cases from Fig. 7 with varying resource supply (*a*_1_, *a*_2_). Indeed, the prediction of competition outcome by this approximation scheme agrees with the full system, where coexistence is attained only when the stabilization factor *b*_*s*_ − *b*_*c*_ is larger than the magnitude of the equalization factor |*e* − *s*|. The grey solid line plotted in the phase diagram shows how the ratio of *a*_11_ + *a*_12_ to *a*_21_ + *a*_22_ affects the equalization and stabilization factors while keeping the total amount of resources *A* = *a*_11_ + *a*_12_ + *a*_21_ + *a*_22_ fixed. For small asymmetries in resource supply (e.g. Fig. 7b), the equalization factor has a small magnitude and the two clusters are very similar in “fitness”. As the asymmetry in the resource supply increases, the the magnitude of the equalization factor |*e*− *s*| increases. Nonetheless, this is accompanied by an increase in the stabilization factor *b*_*s*_ − *b*_*c*_, which allows the system to tolerate the increasing difference in “fitness” in order to remain in the coexistence state. For large asymmetries in resource supply (from Fig. 7a,c), the difference in “fitness” is too large that the stabilization factor due to the interactions is no longer strong enough to sustain coexistence. The same trend is observed with a larger total resource supply *A*. But now that the two clusters are more equalized, the stabilization factor also decreases as shown in the phase diagram.

**FIG. 8:**
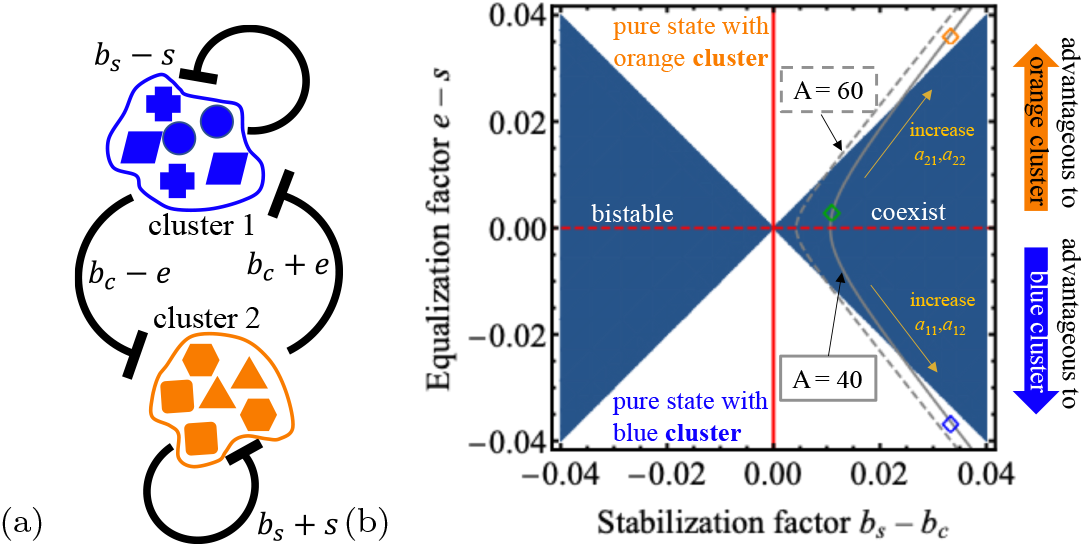
The second level of coarse-graining (Fig. 2b) from taxa to clusters. (a) Graphic representation of Lotka-Volterra approximation, which describes the competition between the two clusters rather than two specialists. (b) Phase diagram showing the three cases from Fig. 7. Total resource supply *A* = Σ_*eℓ*_*a*_*eℓ*_ changes both stabilization and equalization factors. We use a different color from Fig. 6b to show the approximation includes all taxa and does not exclude the generalists.

#### Detection of coarse-graining from timeseries using trend stationarity

Assuming diversity in real systems in the presence of noise is sustained by trade-offs and the dynamics of taxa can be coarse-grained, can we detect clustering using the abundance data? To answer this question, we consider the corresponding Langevin equations (SM Sect. 5) that incorporate noise into the same system of Fig. 7b to generate synthetic data. The time-series generated from the simulation is plotted in Fig. 9a. Is it possible to infer the clustering of taxa into clusters using this set of synthetic data, assuming that we do not know the ground truth? One approach is to consider the trend stationarity of abundances at the cluster level. We define the abundance of cluster *I* as 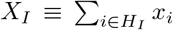, which is plotted in Fig. 9b. Compared to the disorderness due to noise exhibited on the taxon level in Fig. 9a, the cluster abundances are showing mean reversion in the presence of noise: the two clusters coexist at steady state where the mean and variance of the cluster abundances remain constant over time. This corresponds to the first-order autoregression, or AR(1), which is also described as a unit root process and shows a constant trend.

**FIG. 9:**
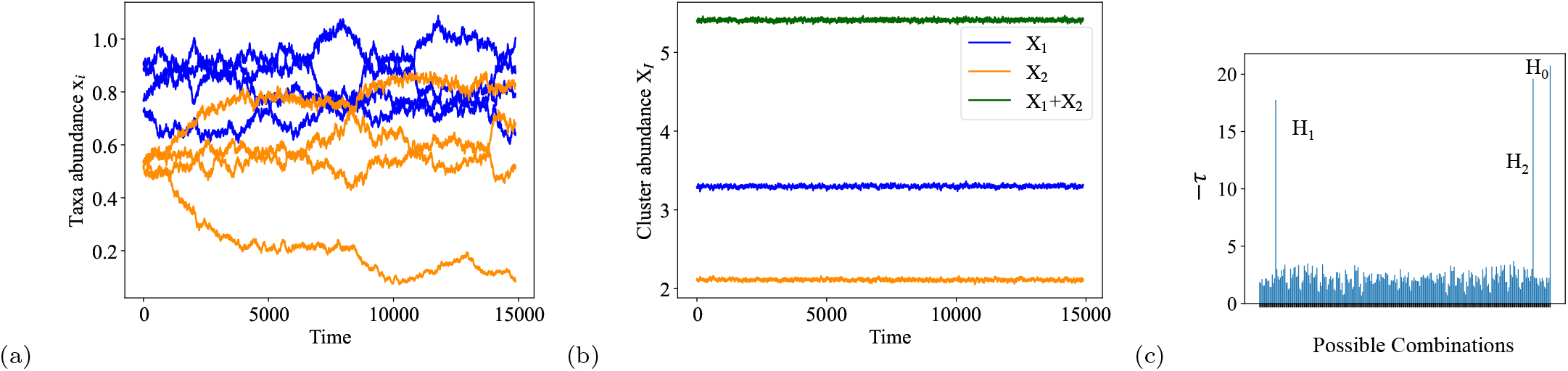
Cluster abundances under second level of coarse-graining (Fig. 2b) exhibit trend stationarity. (a) Abundances of taxa of two different clusters (Blue and orange) in a system exhibiting fluctuations due noise. (b) Cluster abundance *X*_1_, *X*_2_ and the total biomass *X*_0_ shows trend stationarity.(c) Test for trend stationarity of the abundance over all possible combinations of taxa in the powerset *P* (*H*_0_)*\∅* to detect clusters. The three peaks of the test statistics −*τ* correspond to the abundance of the two clusters (*H*_1_, *H*_2_)and the total abundance (*H*_0_).

Suppose we do not know which of the taxa belongs to cluster *I* and given that the system is at steady state on a coarse-grained level, is classification possible through the detection trend stationarity? In other words, given the powerset *P* (*H*_0_) = {{}, {1}, {1, 2}, …, *H*_0_}, which includes all possible ways of combining different taxa, can we find the sets of taxa *H*_*I*_ *⊆ P* (*H*_0_) *\ ∅* (where *∅* = {} is the empty set) that show trend stationarity? To do that, we used the augmented Dickey-Fuller test [62], which is designed as a statistical test for the null hypothesis where the time series is generated by a unit root process, on every member of *P* (*H*_0_)*\ ∅*. Instead of using the test statistics *τ* to reject the null hypothesis, we plotted −*τ* in Fig. 9c to see if there are any signals being picked up. Indeed, three peaks are observed that are well distinguished from other combinations, and they corresponds to *H*_1_, *H*_2_ and *H*_0_ respectively.

While trend stationary allows detection of clusters from taxa, the method is not scalable: for a system with *M* taxa, the cardinality of *P* (*H*_0_) is 2^*M*^. In the next section, we used a different clustering method on larger communities (~ 20 - 100 taxa), which is also applicable here as shown in SM Sect. 5.

### C. Third level of coarse-graining: Further clustering leads to hierarchical structure

So far, we have shown that under coarse-graining as a consequence of the trade-off in Eq. (17), large systems with rich diversity on the taxa level can be interpreted as small systems with a finite number of clusters, limited by the number of effective resources in Fig. 2b. In the previous system, this means there can be no more than two coexisting clusters. Similar to the first level of coarse-graining in Fig. 2a, however, we show in SM Sect. 4 that the upper limit of the number of coexisting clusters can also be lifted if, in addition to Eq. (17), the following trade-off is imposed on all clusters:

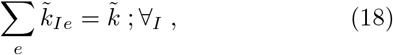

where 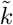 is a constant. Eq. (18) describes the trade-off between the competitiveness of each cluster over essential resources, similar to Eq. (11). This idea of further coarse-graining is illustrated in Fig. 2c. Considering the same system defined by Eq. (8)-Eq. (9) but with the trade-offs Eq. (17) and Eq. (18), is it possible to infer the clustering based on only abundance data when the system size is large?

Specifically, we now consider a system with three clusters *I* = 1, 2, 3, two essential resources *e* = 1, 2, each of which has two alternate sources *N*_*e*_ = 1, 2. Our synthetic data consists of abundance time series 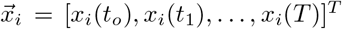 of taxon *i* that still remain in the system at *t* = *T*. Given a dataset of 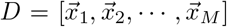 with *M* surviving taxa, the Gram matrix is

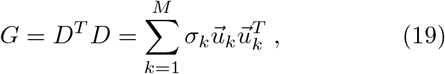

where the last relation is simply the singular value decomposition of *G*, where *k* is assigned based on the singular values such that *σ*_*k*_ *≤ σ*_*k*_*′* for *k > k*′. It turns out that the smallest mode 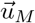 captures fluctuations due to interactions on the cluster level. In Fig. 10a, a large system with three clusters (with a total of thirty taxa) is considered. The matrix 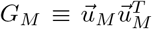 is used for hierarchical clustering, which not only correctly identify the three clusters, namely *H*_1_, *H*_2_ and *H*_3_ but also grouped together clusters with similar competitiveness over essential resources, as shown in Fig. 10b. In SM Sect. 5, we showed the same clustering scheme is also applicable to larger systems with ~100 taxa.

**FIG. 10:**
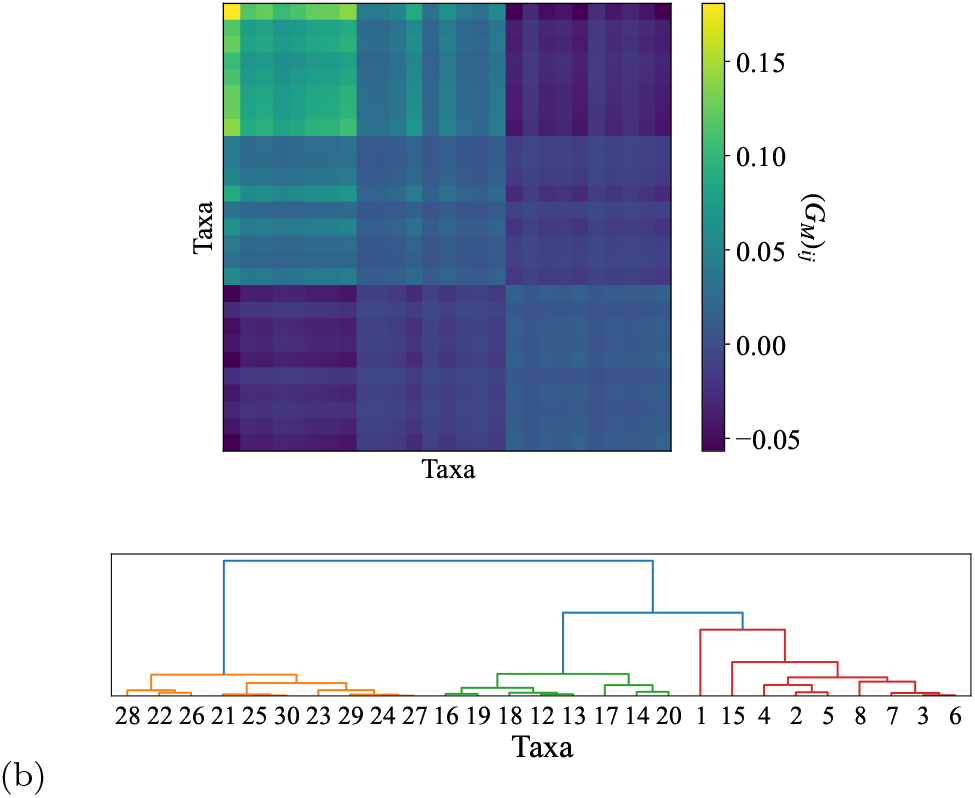
Coarse-graining under the third level of clustering (Fig. 2c) in a system with three clusters from taxa to clusters using Gram matrix. (a) Matrix *G*_*M*_ which corresponds to the smallest singular value of the Gram matrix *G* captures the sub-community structure that allows cluster detection. (b) Hierarchical clustering using matric *G*_*M*_ successfully picked out three clusters *I* = 1, 2, 3, within which taxa share the same set of trade-offs, meaning the same values of 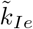 in Eq. (17) and Eq. (18) that corresponds to their competitiveness in essential resources *e*.

## IV. DISCUSSION

Our main result is that hierarchical structures can be sustained through trade-offs. Different from current thinkings of coarse-graining based on similarities, taxa that are clustered share the same trade-off while exhibiting very different phenotypes or nutrient competitiveness. We also propose schemes to detect these strcutures using abundance time series data.

### A. The shielding effect is exhibited beyond competition over alternate resources

Although the idea that trade-offs might have a role in sustaining diversity is not novel [63][54][19][20][22], this is, to our knowledge, the first time that the shielding effect is demonstrated in models beyond competition over alternate resources. In a separate but related paper [11], we showed how toxin-producing taxa can generate the shielding effect to sustain diversity. Here, we demonstrated for the first time that the shielding effect is equally valid for essential resources provided that four conditions are met. First, the population shares the same proteome constraint in these competitions. This means the ZNGIs can be described by smooth functions of the resource levels, as contrasted with the models involving piecewise functions such as the Liebig’s law of minimum [50], and that all taxa share the same maximum growth rate *g*_*m*_. This is traditionally classified as interactive essential resources. Second, there exists a trade-off between a taxon’s competitiveness for the different resources that takes the form of Eq. (11). Third, for each taxon, there exists a trade-off between the competitiveness and stoichiometry for each resource, which is not uncommon for resource competition models involving two strains [49][54]. This is not too dissimilar from the systems with alternate resources, where coexistence requires a trade-off between nutrient efficiency and the impact ratio 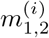 as shown in Eq. (13), which has been implicitly invoked in prior work by setting *S*_*ji*_ = 1 [19][22]. Fourth, the differences in the external supply of the various resources are not too large, so that the consortium is able to self-organize to shield off this asymmetry in resource supply. During shielding, the levels of all resources in this environment sustained by the consortium are at comparable range, allowing all taxa to share the same fitness (or growth rate). This is the first level of coarse-graining as illustrated in Fig. 2a.

To further generalize our framework, we then considered the possibility of generating the shielding effect in systems with a mix of both alternate and essential resources, which has not been previously addressed. Indeed, this is shown to be possible given that taxa within each cluster shares the trade-off in Eq. (17) that gives rise in the second level of coarse-graining (Fig. 2b). We then showed a further coarse-graining (or the third level, in Fig. 2c) if these clusters in the population share the same trade-offs in Eq. (17). In SM Sect. 4, we showed analytically that the third level can be viewed as having one effective single macro-cluster surviving on one effective resource. The implication is that, given the many trade-offs observed in nature [64] that go beyond the competitiveness for alternate resources, it is likely that these other modes of interactions can also sustain shielding effect and contribute to diversity in real systems.

### B. Trade-off promotes diversity by stabilizing the community

How does this structure stabilize the system? To show the role of trade-offs in promoting diversity, we specifically focus on mapping these large systems onto a two-cluster competition, which is the simplest model of competition that can be conceived and has been very well-studied. According to contemporary coexistence theory, a strong intra-species coupling compared to inter-species coupling stabilizes the coexistence state and overcomes the differences in fitness (or equalization factor). The prediction by the theory matches with the behaviour of our full system. Using the Lotka-Volterra competition model to approximate our full system, we showed that this is indeed the case at different levels of coarse-graining such that the system is stabilized by the interactions between the specialists within the same cluster (second level in Fig. 2b) as well as the interactions between sub-communities in a multi-cluster competition (third level in Fig. 2c). In the regime where the system has high diversity, both show negative-frequency selection: strong competition of a taxon or a sub-community with itself implies its fitness decreases when its abundance (or frequency) increases. This provides feedback to keep that taxa or cluster from dominating the population.

### C. Coarse-graining as a path towards diversification

How does diversity develop in microbial communities against competitive exclusion under selective pressure? And how do small systems with low diversity develop into big systems with many strains and species? Trade-offs provide a potential solution by preventing the system from being overdetermined. Symmetry through trade-offs changes the number of effective variables and number of equations to solve, thus allowing large systems to satisfy the competitive exclusion principle, despite diversification (SM Sect. 4). Naturally arising from our analysis is to go beyond one single effective cluster studied in prior work [19][20][22][21] and consider the interactions between multiple clusters. In evaluating the effective diversity in a system, coarse-graining is now possible: taxa that share the same trade-offs can be classified as a cluster. The idea of having some coarse-grained variables is not entirely new: previous clustering schemes were mostly based on similarities between taxa or functional groups [27]. In our case, we show that rather than similarities, it is the variations within each cluster that are desirable: dissimilar taxa specialized at different resources stabilize the cluster and lead to shielding. Within each cluster, these taxa do not share similarities in their strategies on which resource to use; rather, they share a common set of local constraints. As a result, taxa classified as the same cluster share the same fitness, and the competition outcome depends on the difference in fitness between various clusters. One simplistic picture of how early bacterial communities may diversify is as follows: while strong selective pressure keeps the number of coexisting species low, these species can diversify through varying their strategy while upholding a set of trade-offs. This ensures the fitness landscapes remain relatively flat, despite the intra-species variations, which facilitates further speciation when the environment changes.

## ACKNOWLEDGMENTS

We thank members of Prof. Anton Zilman’s group and our group for discussions of ideas. We acknowledge funding from the Natural Sciences and Engineering Research Council of Canada.

## References

[1] G. E. Hutchinson, The Paradox of the Plankton, THE AMERICAN NATURALIST, 9.

[2] R. P. Maharjan, T. Ferenci, P. R. Reeves, Y. Li, B. Liu, and L. Wang, The multiplicity of divergence mechanisms in a single evolving population, Genome Biology 13, 1 (2012).

[3] G. Hardin, The Competitive Exclusion Principle, Science 131, 1292 (1960).

[4] R. A. Armstrong and R. McGehee, Competitive Exclusion, The American Naturalist 115, 151 (1980).

[5] R. McGehee and R. A. Armstrong, Some mathematical problems concerning the ecological principle of competitive exclusion, Journal of Differential Equations 23, 30 (1977).

[6] N. A. Thomopoulos, D. V. Vayenas, and S. Pavlou, On the coexistence of three microbial populations competing for two complementary substrates in con®gurations of interconnected chemostats, Mathematical Biosciences, 16 (1998).

[7] P. Lenas and S. Pavlou, Coexistence of three competing microbial populations in a chemostat with periodically varying dilution rate, Mathematical Biosciences 129, 111 (1995).

[8] J. Huisman and F. J. Weissing, Biodiversity of plankton by species oscillations and chaos, Nature 402, 407 (1999), number: 6760.

[9] C. Lobry and J. Harmand, A new hypothesis to explain the coexistence of n species in the presence of a single resource, Comptes Rendus Biologies 329, 40 (2006).

[10] S. Chakraborty, A. Ramesh, and P. S. Dutta, Toxic phytoplankton as a keystone species in aquatic ecosystems: stable coexistence to biodiversity, Oikos 125, 735 (2016).

[11] G. C. Lui and S. Goyal, Promoting diversity in ecological systems through toxin production, bioRxiv, 2023 (2023).

[12] P. Gerlee and T. Lundh, Productivity and Diversity in a Cross-Feeding Population of Artificial Organisms, Evolution 64, 2716 (2010).

[13] M. J. A. v. Hoek and R. M. H. Merks, Emergence of microbial diversity due to cross-feeding interactions in a spatial model of gut microbial metabolism, BMC Systems Biology 11, 56 (2017).

[14] M. S. Roman and A. Wagner, An enormous potential for niche construction through bacterial cross-feeding in a homogeneous environment, PLOS Computational Biology 14, e1006340 (2018).

[15] R. Marsland III, W. Cui, J. Goldford, A. Sanchez, K. Korolev, and P. Mehta, Available energy fluxes drive a transition in the diversity, stability, and functional structure of microbial communities, PLoS computational biology 15, e1006793 (2019).

[16] P. Mehta and R. Marsland III, Cross-feeding shapes both competition and cooperation in microbial ecosystems (2021), arXiv:2110.04965 [cond-mat, q-bio].

[17] R. Marsland, W. Cui, and P. Mehta, A minimal model for microbial biodiversity can reproduce experimentally observed ecological patterns, Scientific Reports 10, 3308 (2020).

[18] R. Corral López, J. A. Bonachela, M. G. Dominguez-Bello, M. Manhart, S. A. Levin, M. J. Blaser, and M. A. Munõz, Imbalance in gut microbial interactions as a marker of health and disease, Science 391, 890 (2026).

[19] A. Posfai, T. Taillefumier, and N. S. Wingreen, Metabolic Trade-Offs Promote Diversity in a Model Ecosystem, Physical Review Letters 118, 10.1103/PhysRevLett.118.028103 (2017).

[20] T. Taillefumier, A. Posfai, Y. Meir, and N. S. Wingreen, Microbial consortia at steady supply, eLife 6, 10.7554/eLife.22644 (2017).

[21] M. Tikhonov and R. Monasson, Collective Phase in Resource Competition in a Highly Diverse Ecosystem, Physical Review Letters 118, 048103 (2017).

[22] Z. Li, B. Liu, S. H.-J. Li, C. G. King, Z. Gitai, and N. S. Wingreen, Modeling microbial metabolic trade-offs in a chemostat, PLOS Computational Biology 16, e1008156 (2020).

[23] M. Hartmann and J. Six, Soil structure and microbiome functions in agroecosystems, Nature Reviews Earth & Environment 4, 4 (2023).

[24] L. Yan, M. Herrmann, B. Kampe, R. Lehmann, K. U. Totsche, and K. Küsel, Environmental selection shapes the formation of near-surface groundwater microbiomes, Water Research 170, 115341 (2020).

[25] B. Ruth, S. Peter, B. Ibrahim, and P. Dittrich, Revealing the hierarchical structure of microbial communities, Scientific reports 14, 11202 (2024).

[26] Y. Jiang, L. Che, and S. Li, Deciphering the personalized functional redundancy hierarchy in the gut microbiome, Microbiome 10.1186/s40168-025-02273-w (2025).

[27] M. Tikhonov, Theoretical microbial ecology without species, Physical Review E 96, 032410 (2017).

[28] C. Burke, P. Steinberg, D. Rusch, S. Kjelleberg, and T. Thomas, Bacterial community assembly based on functional genes rather than species, Proceedings of the National Academy of Sciences 108, 14288 (2011).

[29] P. J. Turnbaugh, M. Hamady, T. Yatsunenko, B. L. Cantarel, A. Duncan, R. E. Ley, M. L. Sogin, W. J. Jones, B. A. Roe, and J. P. Affourtit, A core gut microbiome in obese and lean twins, nature 457, 480 (2009).

[30] S. Louca, L. W. Parfrey, and M. Doebeli, Decoupling function and taxonomy in the global ocean microbiome, Science 353, 1272 (2016).

[31] S. Louca, M. F. Polz, F. Mazel, M. B. N. Albright, J. A. Huber, M. I. O’Connor, M. Ackermann, A. S. Hahn, D. S. Srivastava, S. A. Crowe, M. Doebeli, and L. W. Parfrey, Function and functional redundancy in microbial systems, Nature Ecology & Evolution 2, 936 (2018).

[32] J. E. Goldford, N. Lu, D. Bajić, S. Estrela, M. Tikhonov, A. Sanchez-Gorostiaga, D. Segrè, P. Mehta, and A. Sanchez, Emergent simplicity in microbial community assembly, Science 361, 469 (2018).

[33] X. Shan, A. Goyal, R. Gregor, and O. X. Cordero, Annotation-free discovery of functional groups in microbial communities, Nature Ecology & Evolution 7, 716 (2023).

[34] Y. Zhao, O. X. Cordero, and M. Tikhonov, Linear-regression-based algorithms can succeed at identifying microbial functional groups despite the nonlinearity of ecological function, preprint (Ecology, 2024).

[35] S. G. Acinas, V. Klepac-Ceraj, D. E. Hunt, C. Pharino, I. Ceraj, D. L. Distel, and M. F. Polz, Fine-scale phylogenetic architecture of a complex bacterial community, Nature 430, 551 (2004), number: 6999.

[36] A. Goyal, L. S. Bittleston, G. E. Leventhal, L. Lu, and O. X. Cordero, Interactions between strains govern the eco-evolutionary dynamics of microbial communities, preprint (Microbiology, 2021).

[37] M. Scott and T. Hwa, Bacterial growth laws and their applications, Current Opinion in Biotechnology 22, 559 (2011).

[38] M. Scott, S. Klumpp, E. M. Mateescu, and T. Hwa, Emergence of robust growth laws from optimal regulation of ribosome synthesis, Molecular Systems Biology 10, 747 (2014).

[39] C. You, H. Okano, S. Hui, Z. Zhang, M. Kim, C. W. Gunderson, Y.-P. Wang, P. Lenz, D. Yan, and T. Hwa, Coordination of bacterial proteome with metabolism by cyclic AMP signalling, Nature 500, 301 (2013).

[40] L. Chao and B. R. Levin, Structured habitats and the evolution of anticompetitor toxins in bacteria., Proceedings of the National Academy of Sciences 78, 6324 (1981).

[41] R. Durrett and S. Levin, Allelopathy in Spatially Distributed Populations, Journal of Theoretical Biology 185, 165 (1997).

[42] L. Pagie and P. Hogeweg, Colicin Diversity: a Result of Eco-evolutionary Dynamics, Journal of Theoretical Biology 196, 251 (1999).

[43] B. Kerr, M. A. Riley, M. W. Feldman, and B. J. M. Bohannan, Local dispersal promotes biodiversity in a real-life game of rock–paper–scissors, Nature 418, 171 (2002).

[44] B. C. Kirkup and M. A. Riley, Antibiotic-mediated antagonism leads to a bacterial game of rock–paper–scissors in vivo, Nature 428, 412 (2004).

[45] M. Whiteley, E. Brown, and R. J. C. McLean, An inexpensive chemostat apparatus for the study of microbial biofilms, Journal of Microbiological Methods 30, 125 (1997).

[46] U.-Y. Pen, C. J. Nunn, and S. Goyal, An Automated Tabletop Continuous Culturing System with Multicolor Fluorescence Monitoring for Microbial Gene Expression and Long-Term Population Dynamics, ACS Synthetic Biology 10, 766 (2021).

[47] M. De Paepe, V. Gaboriau-Routhiau, D. Rainteau, S. Rakotobe, F. Taddei, and N. Cerf-Bensussan, engTrade-off between bile resistance and nutritional competence drives Escherichia coli diversification in the mouse gut, PLoS genetics 7, e1002107 (2011).

[48] J. N. Lester, R. Perry, and A. H. Dadd, Cultivation of a mixed bacterial population of sewage origin in the chemostat, Water Research 13, 545 (1979).

[49] D. Tilman, Resources: A Graphical-Mechanistic Approach to Competition and Predation, The American Naturalist 116, 362 (1980).

[50] Q. Paris, The von Liebig Hypothesis, American Journal of Agricultural Economics 74, 1019 (1992), eprint: https://onlinelibrary.wiley.com/doi/pdf/10.2307/1243200.

[51] L. Pacciani-Mori, S. Suweis, and A. Maritan, Adaptive consumer-resource models can explain diauxic shifts and the violation of the Competitive Exclusion Principle, bioRxiv 10.1101/385724 (2018).

[52] V. Behrends, R. P. Maharjan, B. Ryall, L. Feng, B. Liu, L. Wang, J. G. Bundy, and T. Ferenci, A metabolic trade-off between phosphate and glucose utilization in Escherichia coli, Molecular BioSystems 10, 2820 (2014).

[53] K. F. Edwards, C. A. Klausmeier, and E. Litchman, A Three-Way Trade-Off Maintains Functional Diversity un-der Variable Resource Supply., The American Naturalist 182, 786 (2013).

[54] T. L. S. Vincent, D. Scheel, J. S. Brown, and T. L. Vincent, Trade-Offs and Coexistence in Consumer-Resource Models: It all Depends on what and where you Eat, The American Naturalist 148, 1038 (1996).

[55] M. Mori, E. Marinari, and A. De Martino, A yield-cost tradeoff governs Escherichia coli’s decision between fermentation and respiration in carbon-limited growth, npj Systems Biology and Applications 5, 16 (2019).

[56] S. Dedrick, V. Warrier, K. P. Lemon, and B. Momeni, When does a Lotka-Volterra model represent microbial interactions? Insights from in vitro nasal bacterial communities, mSystems 8, e00757 (2023).

[57] O. S. Venturelli, A. V. Carr, G. Fisher, R. H. Hsu, R. Lau, B. P. Bowen, S. Hromada, T. Northen, and A. P. Arkin, Deciphering microbial interactions in synthetic human gut microbiome communities, Molecular Systems Biology 14, e8157 (2018).

[58] P. H. Kloppers and J. C. Greeff, Lotka–Volterra model parameter estimation using experiential data, Applied Mathematics and Computation 224, 817 (2013).

[59] J. D. Davis, D. V. Olivença, S. P. Brown, and E. O. Voit, EnglishMethods of quantifying interactions among populations using Lotka-Volterra models, Frontiers in Systems Biology 2, 10.3389/fsysb.2022.1021897 (2022).

[60] P. Chesson, Mechanisms of Maintenance of Species Diversity, Annual Review of Ecology and Systematics 31, 343 (2000).

[61] A. D. Letten, P.-J. Ke, and T. Fukami, Linking modern coexistence theory and contemporary niche theory, Ecological Monographs 87, 161 (2017).

[62] S. Seabold and J. Perktold, Statsmodels: Econometric and statistical modeling with python, in Proceedings of the 9th Python in Science Conference, Vol. 57 (Austin, TX, 2010) pp. 10–25080.

[63] D. Tilman, Constraints and Tradeoffs: Toward a Predictive Theory of Competition and Succession, Oikos 58, 3 (1990).

[64] T. Ferenci, Trade-off Mechanisms Shaping the Diversity of Bacteria, Trends in Microbiology 24, 209 (2016).

