## Supplementary Materials for "From shielding effect to hierarchical structures: a coarse-grained description of diversity"

Ga Ching Lui and Sidhartha Goyal

This document presents the supplementary materials for the paper "From shielding effect to hierarchical structures: a coarse-grained description of diversity".

#### 1 Bacterial growth under proteome constraint

Our model is based on [1, 2], which considers a coarse-grained model of protein synthesis limited by metabolism and translation, the rates of which determine the in-flux and out-flux of some intermediates (e.g., amino acids) with level $I$. These rates are dependent on the proteome fractions $\phi_M$ and $\phi_R$, respectively. The exponential growth rate can be written as

$$g = \frac{1}{M}\frac{dM}{dt} = \gamma\phi_R = \nu\phi_M \ , \tag{S.1}$$

where $\gamma$ are $\nu$ are the proportionality constants, which are termed translation efficiency and nutrient efficiency. The last equality in Eq. (S.1) comes from steady state assumption of $\dot{I} = 0$ at the exponential phase. Here, we are describing the growth rate of a single taxon, and for shorthand purposes we did not use the subscript $i$ as in Eq. (1) of the main text the label the taxa: taxa would be characterized by the differences in the efficiencies $\nu$ and $\gamma$.

To satisfy the two growth laws observed under different nutrient conditions and antibiotics levels, it is hypothesized that minimally a three-partition proteome is required:

$$1 - \phi_o = \phi_M + \phi_R \ , \tag{S.2}$$

where $\phi_o$ is the fraction that remains constant and independent of growth rate, while both $\phi_M$ and $\phi_R$ vary. This is known as the proteome constraint Solving for $\phi_R$ in Eq. (S.1) under the constraint in Eq. (S.2) allows us to obtain the growth rate

$$g = \frac{1-\phi_o}{\gamma^{-1} + \nu^{-1}} = \frac{g_m}{R_o + R} \ . \tag{S.3}$$

This is analogous to the circuitry analogy where $g_m$ is the total voltage, and the resistances of the two loads are $R_o$ and $R$. In other words, bacterial growth rate $g_1$ is limited by resource-independent factors characterized by $R_o$ and the resource-dependent factors are characterized by $R$. In the nutrient-limited regime, $R$ is large compared to $R_o$. Our various models in the paper consider different forms of $R$, depending on which modes of resource competitions are involved.

##### 1.1 Growth rate with two alternate resources

For alternate resources, bacterial growth is sustained given that at least one of the resources is present. In the circuitry analogy, this is represented by having two loads connected in parallel, with resistance $R_1$ and $R_2$ respectively. Therefore, the combined resistance is $R^{-1} = R_1^{-1} + R_2^{-1}$: $R$ is large only when both $R_1$ and $R_2$ are large. We will explain this intuition using a coarse-grained model as described below.

In the context of proteome constraint, the influx of $I$ is limited by the proteome fraction $\phi_M$ which includes transporter proteins, and is contributed by both $R_1$ and $R_2$.

$$J_I^{in} \propto \sum_j \tilde{k}_j \eta_j \phi_M = \frac{1}{\beta} \sum_j \tilde{k}_j \eta_j \phi_M \equiv \Sigma_j \nu_j \phi_M \equiv \nu \phi_M \ , \tag{S.4}$$

where the influx $J_I^{in}$ is proportional to $\tilde{k}_j$ and $\eta_j$, which are the efficiency and fraction of the transporter proteins within $\phi_M$ respectively, and $\beta$ is a proportionality constant. In nutrient-limited regime, the efficiency can be modeled as the Monod factor that depends on resource level

$$\tilde{k}_j = k_j^{max} \frac{n_j}{K_j + n_j} \approx k_j^{max} \frac{n_j}{K_j} \ , \tag{S.5}$$

where $k_j^{max}$ is the efficiency when resource $j$ is in abundance and $K_j$ is the Monod constant. The second approximate in Eq. (S.5) is valid when $n_j << K_j$. Therefore $R_j \equiv v_j^{-1} = (\beta K_j / k_j^{max} \eta_j)(1/n_j) \equiv k_j / n_j$. From Eq. (S.1), Eq. (S.4) - (S.5), we recover Eq. (6) in the main text:

$$g = \frac{g^m}{R_o + (n_1/k_1 + n_2/k_2)^{-1}} \ . \tag{S.6}$$

Therefore, the tradeoff in Eq. (10) in the main-text, or in general, $\Sigma_j k_j^{-1} = \tilde{k}^{-1}$ is simply a statement that describes the fractions of proteins used for the alternate resources is fixed: $\Sigma_j \eta_j = 1$.

##### 1.2 Growth rate with two essential resources

For essential resources, we based our derivation on [1, 2, 3], but consider the efficiencies to be dependent on resource level. Here is a quick summary from [3]. For the case of two nutrients $N_1$ and $N_2$, we consider two intermediates $I_1$ and $I_2$ (instead of only having one intermediate $I$), with three sub-processes (1, 2, and R), as shown in Fig. S1.

$$\frac{dI_1}{dt} = \nu_1 \phi_1 - \nu_2 \phi_2 \ ; \tag{S.7}$$

$$\frac{dI_2}{dt} = \nu_2 \phi_2 - \gamma \phi_R \ , \tag{S.8}$$

subjected to the following proteome constraint:

$$1 - \phi_o = \phi_1 + \phi_2 + \phi_R \ . \tag{S.9}$$

The variable $\phi_R$ is the fraction of the proteome that scales with translation, while $\phi_1$ and $\phi_2$ are the metabolic proteins. These fractions vary at different growth rate. Imposing flux balance such that $\dot{I}_1 = \dot{I}_2 = 0$ in Eq. (S.7) and Eq. (S.8) gives the functional form of growth rate

$$g = \frac{1 - \phi_o}{\gamma^{-1} + \nu_1^{-1} + \nu_2^{-1}} \ , \tag{S.10}$$

where $\nu_1$ and $\nu_2$ are the nutrient efficiencies for sub-processes 1 and 2. Similar to before [3], we take $\nu_1$ and $\nu_2$ to be functions of nutrient level $n_1$, $n_2$. Different from the case of alternate resources, the proteome fraction $\phi_M$ is partitioned into $\phi_1$ and $\phi_2$. The trade-off in Eq. (11) of the main-text, or $\Sigma_j k_{ij} = \tilde{k}$, describes a change in the Monod constants $K_j$ in Eq. S.5. Therefore, as efficiency in utilizing one of the resources increases, the species is less efficient at growing on the other resource [4, 5].

#### 2 Derivation of the convex hull and scaling of nutrient levels

While [6] also shows shielding effect in competitions over alternate resources, the functional used for bacterial growth rate is different from our model. In this section, we will use a similar approach from [6] to derive the convex hull applicable to our model for both modes of competition: alternate and essential resources.

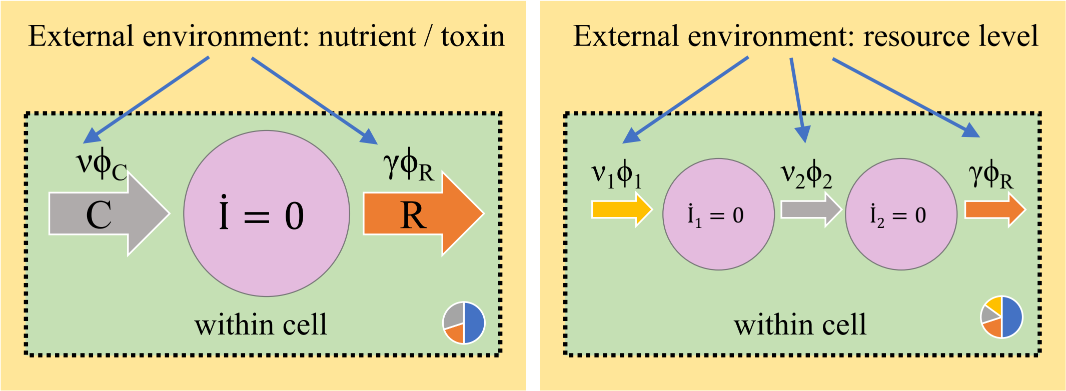


Figure S1: A illustration of a proteome partitioning model. Rate of bacterial growth depends on both proteome partitioning within the cell and abiotic factors in the external environment, for example the level of nutrients and toxins. (Left) Suppose the rate of bacterial growth is limited by two sub-processes C and R, which are responsible for the in-flux and out-flux of an intermediate with abundance I. The rate of the sub-processes is $\nu\phi_C$ and $\gamma\phi_R$, where $\nu$ and $\gamma$ depend on the external environment while $\phi_C$ and $\phi_R$ are the proteome fractions, subjected under the constraint as shown in the inset, in which the blue fraction corresponds to the proteome fraction that is independent of growth rate. (Right) The bacterial growth is limited by three sub-processes involving two intermediates $I_1$ and $I_2$. The efficiencies $v_1$ and $v_2$ depends on the level of different essential resources in the environment, and the proteome fractions $\phi_1$, $\phi_2$ and $\phi_R$ are subjected under the proteome constraint.

#### Alternate resources

For alternative resources, setting $g_i = F$ at equilibrium gives the condition for shielding effect $g_m/F - 1 = \tilde{k}^{-1}/n^\star$. Therefore, we set the scale for nutrient level as

$$L_n = \tilde{k}^{-1}(\frac{g_m}{F} - 1)^{-1} \ . \tag{S.11}$$

To derive the convex hull, notice the sub-current supplied by resource $j$ at shielding is

$$g_{ij} = \frac{g_m n_j/k_{ij}}{1+\Sigma_{j'} n_{j'}/k_{ij'}} = \frac{g_m n^\star/k_{ij}}{1+\Sigma_{j'} n^\star/k_{ij'}} = \frac{g_m \tilde{k}/k_{ij}}{1+\tilde{k}/n^\star} = F\frac{\tilde{k}}{k_{ij}} \ , \tag{S.12}$$

where the last equality comes from $g_i = F$. Setting $\dot{n}_j = 0$ at steady state gives

$$(a_j - n^\star) = \Sigma_i g_{ij} x_i = \tilde{k}\Sigma_i (x_i/k_{ij}) \ . \tag{S.13}$$

Notice that this can be interpreted as the conservation of mass: the total biomass of the cluster (or "species"), which is the weighted sum of the taxa (or "strains") abundances, is contributed by the resources consumed. Consider a change of variables $n_j \to n_j/L_n$, we can write down the convex hull as

$$\{x_i > 0 \quad \forall_i \ ; \quad L_n^{-1}\vec{n}^\star + x_1 L_n^{-1}\tilde{k}\vec{v}_1 + x_2 L_n^{-1}\tilde{k}\vec{v}_2 + \cdots + x_M L_n^{-1}\tilde{k}\vec{v}_M = L_n^{-1}\vec{a}\} \ , \tag{S.14}$$

where $\vec{v}_i = (1/k_{i1}, 1/k_{i2}, \cdots, 1/k_{iN})^T$. For $N = 2$, we can therefore define the impact line as $n_1/L_n = m_i(n_2/L_N - n_2^\star/L_N) + n_1^\star/L_T$ with slope $m_i = k_{i2}/k_{i1}$.

#### Essential resources

For essential resources, setting $g_i = F$ at equilibrium gives the condition for shielding effect $g_m/F - 1 = k/n^\star$. Therefore, we set the scale for nutrient level as

$$L_n = \tilde{k}/(\frac{g_m}{F} - 1) \ , \tag{S.15}$$

which gives $n^\star/L_n = 1$. Consider a change of variables $n_j \to n_j/L_n$, the nutrient dynamics is

$$\dot{n}_j/L_n = (a_j/L_n - n_j/L_n)F - \Sigma_i x_i g_i S_{ji}/L_n \ . \tag{S.16}$$

At steady state $\dot{n}_j = 0$ and $g_i = F$, the supply concentration has to be large enough to support coexistence, meaning

$$L_n^{-1}\vec{a} = L_n^{-1}\vec{n}^\star + L_n^{-1}\boldsymbol{S}\vec{x} \; , \tag{S.17}$$

where $\boldsymbol{S}$ is the matrix with elements $S_{ji}$. Feasibility of the coexistence solution simply refers to non-negative abundance for all taxa. This means the supply vector has to lie within the convex hull defined by the stoichiometries $\vec{S}_i = (S_{1i}, S_{2i}, \cdots, S_{Ni})^T$

$$\{x_i > 0 \quad \forall_i \; ; \quad L_n^{-1}\vec{n}^\star + x_1 L_n^{-1}\vec{S}_1 + x_2 L_n^{-1}\vec{S}_2 + \cdots + x_M L_n^{-1}\vec{S}_M = L_n^{-1}\vec{a}\} \; . \tag{S.18}$$

For $N = 2$, we can therefore define the impact line as $n_1/L_n = m_i(n_2/L_N - n_2^\star/L_N) + n_1^\star/L_T$ with slope $m_i = S_{1i}/S_{2i}$.

### 3 Derivation for Lotka-Volterra approximation

In this section, we show our derivation of approximating the full system defined by Eq. (1) in the main text with competitive Lotka-Volterra equations. While [7] takes a similar approach, it is a little trickier to approximate our full model since the nutrient dynamics $\dot{n}$ explicitly depends on growth rate $g_i$. s The general approach involves three steps: (1) assumes separation of timescale by setting $\dot{x} = 0$, such that the dynamics of taxa abundance is faster than the dynamics of nutrients; (2) takes $\dot{n_\ell} = 0$ and solve for nutrient resources as $n_\ell = n_\ell(\vec{x})$ and; (3) compares the ZNGIs with the nullclines of the Lotka-volterra equations:

$$1 - \sum_j b_{ij} x_j = 0 \; . \tag{S.19}$$

Here, we give a short derivation of how to obtain the interaction matrix $B$ for competitions over alternate resources, essential resources, and a combination of both.

#### 3.1 Effective coupling due to alternate resource

The ZNGI for taxon $i$ is given by setting $\dot{x}_i = 0$ in Eq.(1) of the main text, or equivalently, $g_i = F$ where the growth rate is given by Eq. (6):

$$\left(\sum_\ell \frac{n_\ell}{k_{i\ell}}\right)^{-1} = \frac{g_m}{F} - 1 \equiv G \; . \tag{S.20}$$

We proceed to taking $\dot{n} = 0$ in Eq. (1). Substituting Eq. (S.20) into the equation gives

$$n_\ell = a_\ell - \sum_i \frac{S_{\ell i}}{F} \frac{\frac{n_\ell}{k_{i\ell}}}{1 + G^{-1}} g_m x_i \; , \tag{S.21}$$

which can be used to solve for the resource levels

$$n_\ell = a_\ell \left(1 + \sum_i \frac{S_{\ell i}}{k_{i\ell}} G x_i\right)^{-1} \approx a_\ell \left(1 - \sum_i \frac{S_{\ell i}}{k_{i\ell}} G x_i\right) \; . \tag{S.22}$$

The last term comes from the assumption that $x_i$ is very small compared to $k_{i\ell}/GS_{\ell i}$. For each resource $\ell$, we multiply Eq. (S.22) by $1/k_{i\ell}$ and sum up the term across $\ell$. Substitute this into Eq. (S.20), we now have

$$\frac{1}{G} = \left(\sum_\ell \frac{a_\ell}{k_{i\ell}}\right) - \sum_j \left(\sum_\ell \frac{a_\ell S_{\ell j}}{k_{i\ell} k_{j\ell}}\right) G x_j \; . \tag{S.23}$$

Comparing this set of equations with the nullclines in a Lotka-volterra system in Eq. (S.19), the matrix elements of the interaction matrix is

$$b_{ij} = \frac{\left(\sum_\ell \frac{a_\ell S_{\ell j}}{k_{i\ell} k_{j\ell}}\right) G}{\left(\sum_\ell \frac{a_\ell}{k_{i\ell}}\right) - \frac{1}{G}} \; . \tag{S.24}$$

Similar to the case of essential resources in Fig. 5 of the main-text, we compare the Lotka-volterra approximation with competition outcome in the full system shown in Fig. 4c-e for the alternate resources. Using Eq. (S.19) and Eq. (S.24), we showed in Fig. S2 that our approximation gives the same qualitative outcomes as the supply rate $(a_1, a_2)$ changes. These three cases are plotted in the phase diagram in Fig. 6b of the main text.

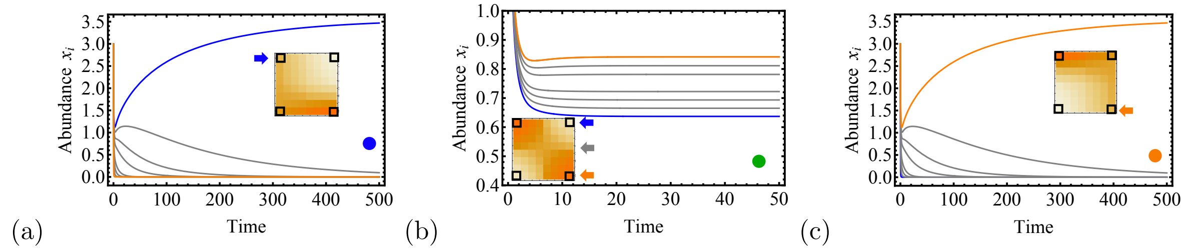


Figure S2: Competitions in the first level of corse-graining are stabilized by specialists for alternate resources. (a-c) Time series of taxa abundances obtained by Lotka-Volterra approximation at three different resource supplies as denoted in Fig. 4a, showing qualitative agreements with full system in Fig. 6c-e. Inset shows the interaction matrix $B_{ij}$, with the color values indicating the coupling strength and the arrow shows the rows correspond to winning taxa. Using the matrix elements at the corners (in black square), we model this as a competition between the two specialists (blue and orange) while ignoring the generalists (gray), with four possible outcomes.

#### 3.2 Effective coupling due to essential resource

At $\dot{x}_i = 0$, the growth rate of the taxa that still survive matches with the flow rate of the chemostat $g_i = F$, where the growth rate is given by Eq. (3). Setting $\dot{n}_\ell$ in Eq. (1) and solving for $n_\ell$ give

$$\frac{1}{n_\ell} = a_\ell^{-1}\left(1 - \sum_i \frac{S_{\ell i}}{a_\ell} x_i\right)^{-1} \approx \frac{1}{a_\ell}\left(1 + \sum_i \frac{S_{\ell i}}{a_\ell} x_i\right) , \tag{S.25}$$

where the last term assumes $x_i << \frac{a_\ell}{S_{\ell i}}$. By multiplying $k_{i\ell}$ to Eq. (S.25) and sum over all $\ell$, we can recover the ZNGI for taxon $i$:

$$G \equiv \frac{g_m}{F} - 1 = \sum_\ell \frac{k_{i\ell}}{n_\ell} = \sum_\ell \frac{k_{i\ell}}{a_\ell} + \sum_j \left(\sum_\ell \frac{k_{i\ell} S_{\ell j}}{a_\ell^2}\right) x_j \, . \tag{S.26}$$

Compare this expression with the nullclines of a Lotka-Volterra system in Eq. (S.19), we obtain the interaction matrix

$$b_{ij} = \frac{\sum_{\ell=1}^{N} \frac{k_{i\ell} S_{\ell j}}{a_\ell^2}}{G - \sum_{\ell=1}^{N} \frac{k_{i\ell}}{a_\ell}} \tag{S.27}$$

#### 3.3 A two-cluster competition

For a $2 \times 2$ interaction matrix $B$, coexistence cooresponds to the $(x_1^\star, x_2^\star)$ of the null-clines:

$$B\mathbf{x} = \begin{pmatrix} b_{11} & b_{12} \\ b_{21} & b_{22} \end{pmatrix} \begin{pmatrix} x_1^\star \\ x_2^\star \end{pmatrix} = \begin{pmatrix} 1 \\ 1 \end{pmatrix} , \tag{S.28}$$

which can be solved using Cramer's rule:

$$\frac{1}{\det(B)} \begin{pmatrix} b_{22} & -b_{12} \\ -b_{21} & b_{11} \end{pmatrix} \begin{pmatrix} 1 \\ 1 \end{pmatrix} = \begin{pmatrix} x_1^\star \\ x_2^\star \end{pmatrix} . \tag{S.29}$$

Therefore, for the coexistence solution to be feasible, namely $x_1^\star, x_2^\star > 0$, the following condition has to be satisfied:

$$\mathrm{sgn}(\det B) = \mathrm{sgn}(b_{22} - b_{12}) = \mathrm{sgn}(b_{11} - b_{21}) , \tag{S.30}$$

where sgn is the signum function. The coexistence state is stable when the determinant of the interaction matrix $\det(B) > 0$. In Eq. (16) of the main-text, we defined the self-interaction strength $b_s = (b_{11} + b_{22})/2$, cross-interaction strength $b_c = (b_{12} + b_{21})/2$, and the two symmetry-breaking terms $s = (b_{22} - b_{11})/2$ and $e = (b_{12} - b_{21})/2$.

It turns out a two-cluster competition in the framework of competitive Lotka-Volterra dynamics can be described purely by the stabilization factor $b_s - b_c$ and equalization factor. Eq. S.30 is simply

$$\mathrm{sgn}(\det B) = \mathrm{sgn}((b_s - b_c) - (e - s)) = \mathrm{sgn}((b_s - b_c) + (e - s)) \ . \tag{S.31}$$

While the determinat involves quadratic terms of $b_s$, $b_c$, $e$ and $s$, the last equality involves only linear terms. Therefore, a stable coexistence solution required the following condition: $b_s - b_c > |e - s|$, which corresponds to Zone (2) in Fig. 6 of the main-text. The self-interaction strength $b_s$ is stronger than the cross-interaction strenghth $b_c$, which stabilizes the coexistence of the two clusters.

Other possible competition outcomes are notated in Fig. 6 as Zone (1), (3) and (4). Zone(3) corresponds to a bistability, such that the coexistence solution $(x_1^\star, x_2^\star)$ is still feasible but unstable: $-\mathrm{sgn}((b_c - b_s) + (e - s)) = -\mathrm{sgn}((b_c - b_s) - (e - s))$, with $b_c > b_s$ and $det(B) < 0$. Zone (1) and (4) corresponds to pure states with either one of the two clusters dominating, since the coexistence solution $(x_1^\star, x_2^\star)$ is no longer feasible: $|e - s| > |b_s - b_c|$.

### 4 Coarse-grainability due to trade-offs

#### 4.1 Coarse-graining in mixed mode of resources

In Sect. III B of the main-text, the model corresponds to a mix of both essential and alternative resources is considered (Eq. (8) - Eq. (9) in the main-text). The bacterial growth rate takes the following form:

$$g_i = \frac{g_m}{1 + \sum_{e=1}^{E} \left( \sum_{\ell=1}^{N_e} \frac{n_{e\ell}}{k_{ie\ell}} \right)^{-1}} \ , \tag{S.32}$$

for which there are $E$ essential nutrients, each has $N_e$ alternative channels that taxon $i$ grows on. At this microscopic level, there are in total $N = \Sigma_e N_e$ number of abiotic factors. Coexistence at the steady state implies the same growth rate for every taxon in the system that balances with the flow rate of the chemostat. Suppose there are $p$ taxa with $i = 1, \ldots, p$, solving the ZNGIs ($g_i = F$) gives a total of $p$ constraints over $N$ variables. Therefore, it is a pathological case to reach coexistence at the steady state if $p > N$, which is known as the competitive exclusion principle.

This can be circumvented by applying constraints, which reduces the number of effective abiotic factors, or in our language the coarse-grained abiotic factors. Consider for each taxon $i \in I$, where $I = 1, 2, \ldots, P$ notates clusters, each satisfying the following $E$ constraints:

$$\sum_\ell \frac{1}{k_{ie\ell}} = \frac{1}{\tilde{k}_{Ie}} \ , \qquad e = 1, 2, \ldots, E \ ; \tag{S.33}$$

where $e = 1, 2, \ldots, E$. These constraints are represented by the blue dotted squares in Fig. 3e. In other words, for each essential resource, we have imposed a local constraint over its alternative channels. The strains can all reach the same growth rate at steady state if $n_{e\ell} = n_e^\star$ for all $\ell$, which is the shielding effect that we have previously discussed. It follows that the growth rate is

$$g_i = \frac{g_m}{1 + \sum_{e=1}^{E} \frac{\tilde{k}_{Ie}}{n_e^\star}} \ , \quad \forall i \in I \ . \tag{S.34}$$

Therefore, the number of effective abiotic factors is changed from $N$ to $E < N$. By the competitive exclusion principle, at steady state there can be at most $P = E$ coexisting clusters, each violating the principle locally with no upper limit imposed on the the number of strains for each clustered group. Intuitively, we have reduced the size of our model from $p$ strains and $N$ abiotic factors to a smaller system with $P$ clusters competing over $E$ essential nutrients. This is the second-level of coarse-graining, as illustrated in Fig. 2b of the main-text. The special that has been explored [ref] is when $E = P = 1$, which is the first level of coarse-graining shown in Fig. 2a of the main-text.

For the third level of coarse-graining (Fig. 2c of the main-text), we consider further coarse-grained by applying the following set of constraints:

$$\sum_e \tilde{k}_{Ie} = K_{I'} \ , \tag{S.35}$$

with $I' = 1, \ldots, P'$. Effectively, this allows further grouping of $P$ clusters into $P' < P$ "macro-clusters", and within each macro-cluster, there is no upper limit imposed on the number of coexisting clusters. In order words, by using different sets of constraints, we can construct a system with a hierarchy of diversity that satisfies the competitive exclusion principle at each level. As a special case, we can apply the same constraint to all clusters

$$K_I = K \ ,$$

then at the steady state, the growth rate of all clustered-groups is simply

$$g_I = \frac{g_m}{1 + \frac{K}{n^\star}} \ ,$$

which has the same form as the equation for a single strain growing on a single growth-limiting resource.

#### 4.2 Coarse-graining in chemostats at steady state

Consider the second level of coarse-graining (Eq. (S.34)), and suppose there is a separation of timescale where the abiotic factors have reached steady state such that $n_{e\ell} = n_e^\star$ for all $\ell$, and $g_i = g_I$ for all strains $i \in I$ as shown in Eq.(S.34), we sum up the abundances of strains of the same cluster and the levels of alternate sources of the same essential nutrient in Eq. (9), which gives

$$\sum_{i \in I} \dot{x}_i = (g_I - F) \sum_{i \in I} x_i \tag{S.36}$$

$$\sum_{\ell=1}^{N_e} \dot{n}_{e\ell} = \left( \sum_{\ell=1}^{N_e} a_{e\ell} - N_e n_e^\star \right) F - \sum_I \left( S_{eI} g_I \sum_{i \in I} x_i \right) \ . \tag{S.37}$$

Thus, Eq. (9) in the main-text is reduced to a lower dimensional description that involves clusters and coarse-grained resources.

$$\dot{X}_I = (g_I - F) X_I \ , \quad I = 1, \ldots, P \ ; \tag{S.38}$$

$$\dot{n}_e = N_e \left( \bar{a}_e - n_e^\star \right) F - \sum_I \left( S_{eI} g_I X_I \right) \ , \quad e = 1, \ldots, E \ , \tag{S.39}$$

with $\bar{a}_e = \Sigma_\ell a_{e\ell} / N_e$ as the average supply concentration across alternate resources, $X_I = \Sigma_{i \in I} x_i$ as the abundance of cluster $I$ and $n_e = \Sigma_\ell n_{e\ell}$. In short summary, by evoking these local constraints, both the taxa and resources can be grouped into coarse-grained variables. It provides one of the plausible mechanisms that resolves the paradox of how to maintain diversity while upholding the competitive exclusion principle: the principle is seemingly violated only on the taxa level, but not on the cluster level. At shielding, the system consists of solving $P$ equations ($g_I = F$) over $E$ variables ($n_e^\star$). Thus, the number of coarse-grained "species", namely the clusters, cannot exceed the number of coarse-grained resources.

### 5 Stochastic simulation

Our treatment is similar to Khatri, Free and Allen (2012) [8]. Suppose we consider $e = 1, 2$ essential nutrients, and each has alternative resources, which are denoted by $\ell, \ell' = 1, 2$, and consider $i = 1, ..., M$ strains. For this system, we consider the following chemical reactions:

$$X_i \xrightarrow{a'_i} \phi \ ; \tag{S.40}$$

$$\phi \xrightarrow{a'_{M+2(e-1)+\ell}} N_{e\ell} \ ; \tag{S.41}$$

$$N_{e\ell} \xrightarrow{a'_{M+4+2(e-1)+\ell}} \phi \; ; \tag{S.42}$$

$$X_i + S_{1i}N_{1\ell} + S_{2i}N_{2\ell'} \xrightarrow{a'_{M+8+4(i-1)+2(\ell-1)+\ell'}} 2X_i \; , \tag{S.43}$$

where the capital letters indicate the chemical species that is involved. The propensity functions $a'_k$ gives the rate of the $k$-thchemical reaction. We also define the stoichiometry vector $\vec{r}_k$, which is a (M+4)-dimensional vector. We notate each element by $(\vec{r}_k)_j$, with $j = 1, 2, ..., M+4$. The elements correspond to the stoichiometries of reaction $k$ to $(X_1, X_2, ..., X_M, N_{11}, N_{12}, N_{21}, N_{22})$. Unless otherwise specified below, the values of $(\vec{r}_k)_j$ are set to be 0.

$$a'_i = Fx_i \; ; \qquad (\vec{r}_i)_i = -1 \tag{S.44}$$

$$a'_{M+2(e-1)+\ell} = a_{e\ell}F \; ; \qquad (\vec{r}_{M+2(e-1)+\ell})_{(M+2(e-1)+\ell)} = 1 \tag{S.45}$$

$$a'_{M+4+2(e-1)+\ell} = Fn_{e\ell} \; ; \qquad (\vec{r}_{M+4+2(e-1)+\ell})_{M+2(e-1)+\ell} = -1 \tag{S.46}$$

$$a'_{M+8+4(i-1)+2(\ell-1)+\ell'} = g_i \frac{g_{i1\ell}}{g_i}\frac{g_{i2\ell'}}{g_i} x_i \; ; (\vec{r}_{M+8+4(i-1)+2(\ell-1)+\ell'})_i = 1 \; ; \tag{S.47}$$

$$(\vec{r}_{M+8+4(i-1)+2(\ell-1)+\ell'})_{M+\ell} = -S_{1i}; (\vec{r}_{ijk})_{M+2+\ell'} = -S_2 \; . \tag{S.48}$$

We then use the Euler scheme to solve the following Langevin equation:

$$\frac{d\vec{n}}{dt} = \vec{A}(\vec{n}) + B^{1/2}\vec{\xi}(t) \; , \tag{S.49}$$

where

$$A = \sum_k a'_k \vec{r}_k \; , \tag{S.50}$$

which recovers our deterministic system and the noise is encoded in

$$B = \sum_k a'_k \vec{r}_k \cdot \vec{r}_k^{\,T} \; , \tag{S.51}$$

where $< \xi >= 0$ and $< \xi\xi' >= I\delta(t-t')/V$, with $I$ being the identity matrix and $1/\sqrt{V}$ determines the magnitude of noise. For each step of the numerical scheme, $B^{1/2}$ is obtained by updating vector $a$.

Using the procedure described in Sect. III C of the main text, we use hierarchical clustering on larger systems for both the second and the third level of clustering.

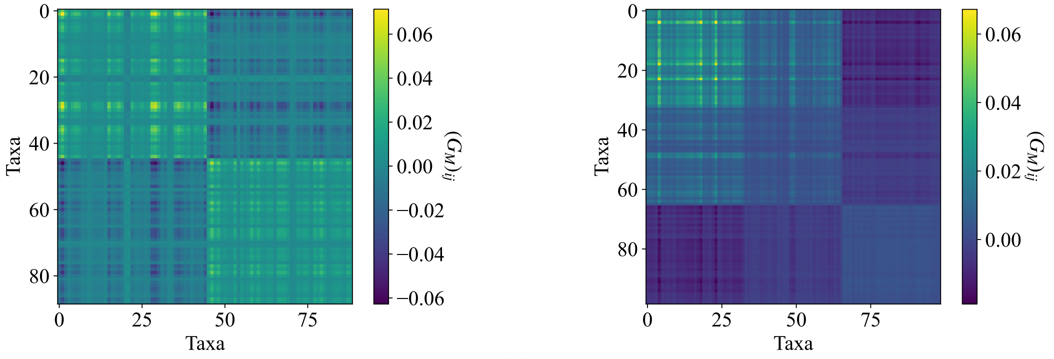


(a) (b)

Figure S3: Coarse-graining for large systems. The Gram matrix from strain abundances show subcommunity structures for both (a) second level and (b) third level of clustering, competing for effectively two essential nutrients.

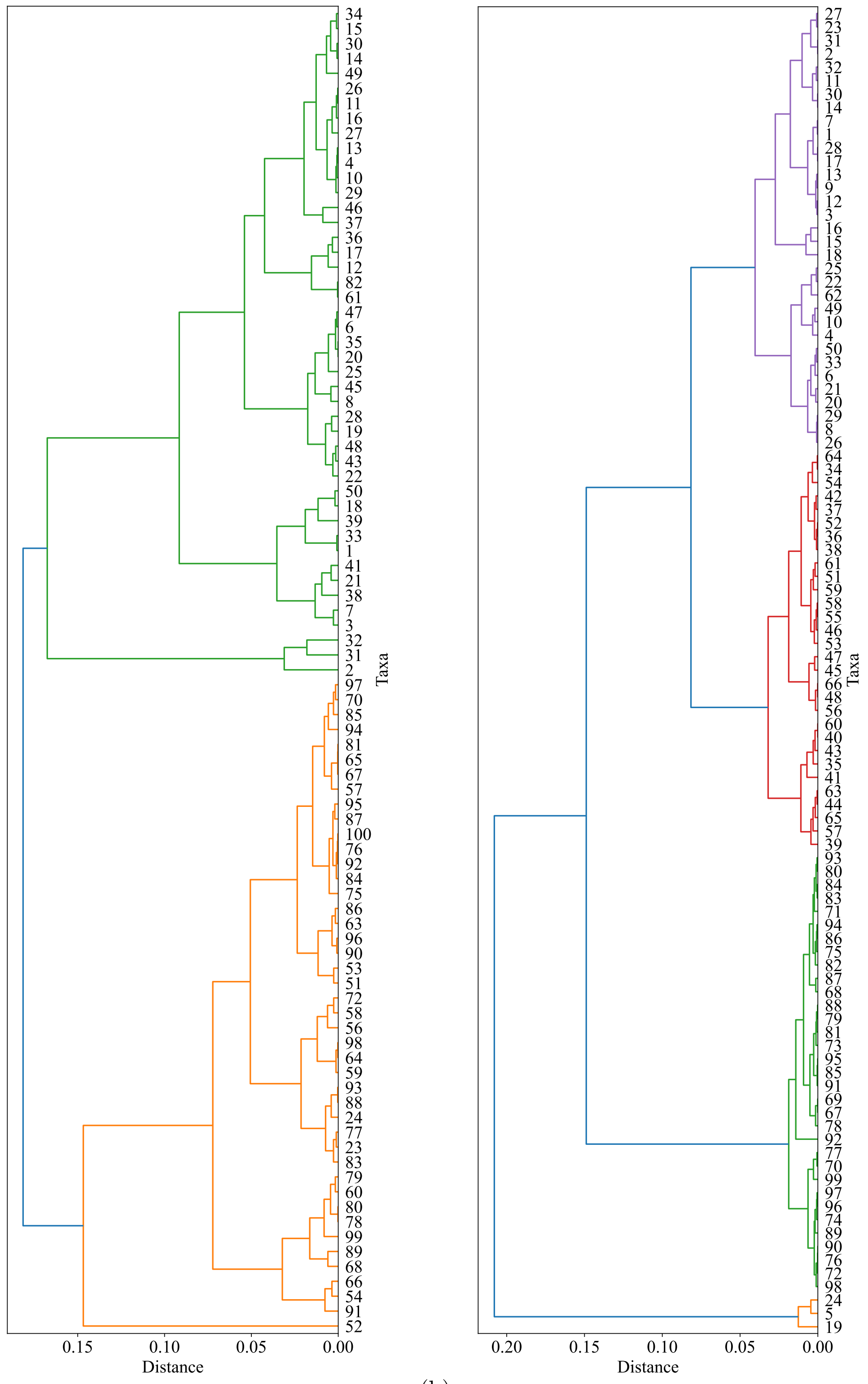


(a) (b)

Figure S4: Hierarchical clustering for large systems based on the Gram matrices in Fig. S3, showing (a) two and (b) three clusters respectively.

### References


[1] C. You, H. Okano, S. Hui, Z. Zhang, M. Kim, C. W. Gunderson, Y.-P. Wang, P. Lenz, D. Yan, and T. Hwa, Nature **500**, 301 (2013).

[2] M. Scott, S. Klumpp, E. M. Mateescu, and T. Hwa, Molecular Systems Biology **10**, 747 (2014).

[3] G. C. Lui and S. Goyal, bioRxiv , 2023.03.06.531399 (2025).

[4] V. Behrends, R. P. Maharjan, B. Ryall, L. Feng, B. Liu, L. Wang, J. G. Bundy, and T. Ferenci, Molecular BioSystems **10**, 2820 (2014).

[5] K. F. Edwards, C. A. Klausmeier, and E. Litchman, The American Naturalist **182**, 786 (2013).

[6] A. Posfai, T. Taillefumier, and N. S. Wingreen, Physical Review Letters **118**, 028103 (2017).

[7] A. D. Letten, P.-J. Ke, and T. Fukami, Ecological Monographs **87**, 161 (2017), _eprint: https://esajournals.onlinelibrary.wiley.com/doi/pdf/10.1002/ecm.1242.

[8] B. S. Khatri, A. Free, and R. J. Allen, Journal of Theoretical Biology **314**, 120 (2012).
